# PI3K-AKT activation determines oncogenic RAS-induced hypertranscription and replication stress

**DOI:** 10.64898/2026.03.16.711577

**Authors:** Richard D. W. Kelly, Claire Wilson, Claire H. M. Tang, Rosanna J. Wilkins, Aditi Kanhere, Eva Petermann

**Affiliations:** Department of Cancer and Genomic Sciences, College of Medicine and Health, University of Birmingham, Birmingham, UK; Birmingham Centre for Genome Biology, University of Birmingham, Birmingham, UK; Department of Molecular & Clinical Cancer Medicine, Institute of Systems, Molecular and Integrative Biology, University of Liverpool, United Kingdom; Division of Medical Sciences, Chester Medical School, University of Chester, UK; Chromosome & Cellular Dynamics Section, Institute of Medical Sciences, University of Aberdeen, Aberdeen, UK; Department of Pharmacology and Therapeutics, Institute of Systems, Molecular & Integrative Biology, University of Liverpool, UK

**Author notes:** Correspondence: Eva Petermann.

**Keywords:** Oncogene, HRAS, KRAS, BRAF, E2F, MYC, transcription-replication conflicts, R-loops, DNA damage, cancer, cell cycle

## Abstract

Hypertranscription and transcription-replication conflicts (TRCs) are frequent features of cancer cells. RAS oncogenes promote hypertranscription to allow cell growth and proliferation, which can the lead to TRCs. Here, we report that hyperactivation of the PI3K-AKT signalling pathway is required for TRCs induced by RAS oncogenes. Oncogenic HRAS causes more TRCs than oncogenic KRAS or BRAF, because HRAS hyperactivates PI3K. PI3K hyperactivation is associated with in glycogen synthase kinase-3β (GSK3β) inhibition, increased E2F and MYC transcription programmes, increased nascent transcription of ribosome biogenesis genes and small nucleolar RNAs (snoRNA) expression. Small molecule inhibition of PI3K signalling prevents RAS-induced replication stress, and small molecule PI3K activation promotes replication stress. RAS-induced TRCs require a cooperation of MAPK and Pi3K signalling, S phase entry and hypertranscription. Our findings suggest a mechanistic explanation for replication stress variability between RAS activation models and identify PI3K pathway activation as a potential new determinant of TRCs in cancer.

## INTRODUCTION

Oncogenic RAS signalling drives cancer cell proliferation through accelerating the cell cycle and activating protein synthesis^1, 2^. RAS proteins are GTPases that, in response to extracellular growth signals, drive proliferation through activation of downstream phosphorylation cascades such as the RAF-MEK-ERK MAP kinase (MAPK) pathway^3^. Oncogenic mutations constitutively activate RAS GTPase activity independently of external growth signals, leading to aberrant proliferation. Three different RAS genes *HRAS*, *KRAS* and *NRAS* encode four RAS isoforms HRAS, KRAS4A, KRAS4B and NRAS, which display different post-translational modifications and subcellular localisation^3, 4^. RAS gene mutations, predominantly in *KRAS*, occur in 10-15% of human cancers, especially in gastrointestinal cancers, lung cancer and multiple myeloma^4, 5, 6^.

De-regulated cell proliferation can promote DNA replication stress^7, 8, 9, 10^, a source of chromosomal instability (CIN)^11^, which can in turn promote aggressiveness and therapy resistance^12^. RAS oncogenes and their downstream effectors such as MYC and CYCLIN E can cause replication stress through increasing replication initiation, transcription activity or cohesin occupancy on chromatin^7, 8, 9, 10^. Oncogene-induced widespread increases in transcription, also called hypertranscription, are proposed to increase the chance of transcription-replication conflicts (TRCs), where the transcription machinery or co-transcriptional RNA-DNA hybrids (R-loops) interfere with replication fork progression^13^. Perturbed replication fork progression can result in accumulation of under-replicated DNA or DNA breaks which in turn promote mitotic aberrations and CIN^11^.

Oncogenic HRAS can cause TRCs through upregulating mRNA and protein levels of the general transcription factor TATA-box binding protein (TBP), leading to increased nascent RNA synthesis (hypertranscription) and RNA:DNA hybrid levels^8^. Oncogenic transcription factors such as YAP1 and TAZ^14^, which are downstream effectors of the HIPPO pathway, and oncogenic TAZ-CAMTA1^15^ and EWS-FLI^16^ transcription factor fusions are also reported to induce hypertranscription, RNA:DNA hybrid accumulation and TRCs. Hypertranscription has been reported across multiple cancer types, with implications for prognosis^17, 18^, and small molecule inhibitors targeting the interaction between replication and transcription are under preclinical investigation for exploiting TRCs in RAS-mutant cancer models^19, 20^. RAS pathway mutations are therefore potential biomarkers for hypertranscription and replication stress in cancer, which could inform prognosis^17, 18^ and therapeutic approaches^20, 21^.

Downstream of RAS, the MAPK pathway can activate RNA polymerases and RNA synthesis for cell growth, with MEK and ERK activity promoting the expression and activity of crucial transcription factors such as MYC, TBP, and the RNA polymerase I transcription factor RRN3^22, 23, 24^. In addition, the PI3K-AKT pathway can act both alongside and downstream of RAS and promote RNA synthesis for cell growth^25^. Both MAPK and PI3K pathway inhibitors are in development for targeted therapy of RAS-mutant cancers^26, 27^. Furthermore, the development of RAS inhibitors has shown that the different signalling properties of specific RAS mutants are important for rational therapeutic targeting^28^.

Understanding the role of MAPK and PI3K signalling pathways in RAS-induced hypertranscription and replication stress will therefore be important to decipher the origins of genomic instability in cancer and help predict oncogene-induced replication stress for precision therapeutic targeting.

Here we investigate how RAS isoforms and key RAS downstream signalling pathways promote hypertranscription and replication stress. Unexpectedly, activation of MAPK signalling alone was insufficient for inducing hypertranscription or replication stress. Instead, our data support that PI3K-AKT signalling is needed for RAS-induced hypertranscription and replication stress, due to the role of the PI3K pathway in promoting RNA synthesis and cell growth. Due to its stronger activation of PI3K signalling, mutant HRAS induces more hypertranscription and replication stress than mutant KRAS. Small molecule inhibition or activation of PI3K can modulate replication stress, and combining PI3K activation with KRAS or BRAF activation can exacerbate replication stress compared to activation of each oncogene alone. Our data suggest that PI3K pathway activation may be a key effector of TRCs, shedding light on determinants of oncogene-induced replication stress.

## RESULTS

### Oncogenic HRAS, KRAS and BRAF induce different levels of hypertranscription and replication stress

To investigate the contribution of RAS downstream pathways to oncogene-induced replication stress, we generated immortalised human BJ fibroblasts expressing tamoxifen-inducible KRAS^G12V^ or doxycycline-inducible BRAF^V600E^. We then used these alongside BJ fibroblasts expressing tamoxifen-inducible HRAS^G12V^, which we had characterised before^8^. Tamoxifen- or doxycycline-induced accumulation of mutant HRAS, KRAS or BRAF proteins hyperactivated the MAP kinase signalling pathway, as evidenced by increased ERK1/2 phosphorylation in all three cell lines (Fig. 1A, B). We then quantified nuclear incorporation of 5-Ethynyluridine (EU) to assess nascent RNA synthesis as a measure of hypertranscription (Fig. 1C). As reported before, nascent RNA synthesis increased up to two-fold after HRAS^G12V^ induction^8^. Unexpectedly however, nascent RNA synthesis increased but only up to 1.2-fold after KRAS^G12V^ induction and not at all after BRAF^V600E^ induction (Fig. 1C, D). We also used S9.6 antibody^29^ slot blots to measure global RNA:DNA hybrid levels in response to oncogene induction. RNase H treatment confirmed specific detection of RNA:DNA hybrids (Fig. 1E). While HRAS^G12V^ induction increased RNA:DNA hybrid levels, as reported before^8^, KRAS^G12V^ or BRAF^V600E^ induction did not.

**Figure 1.**
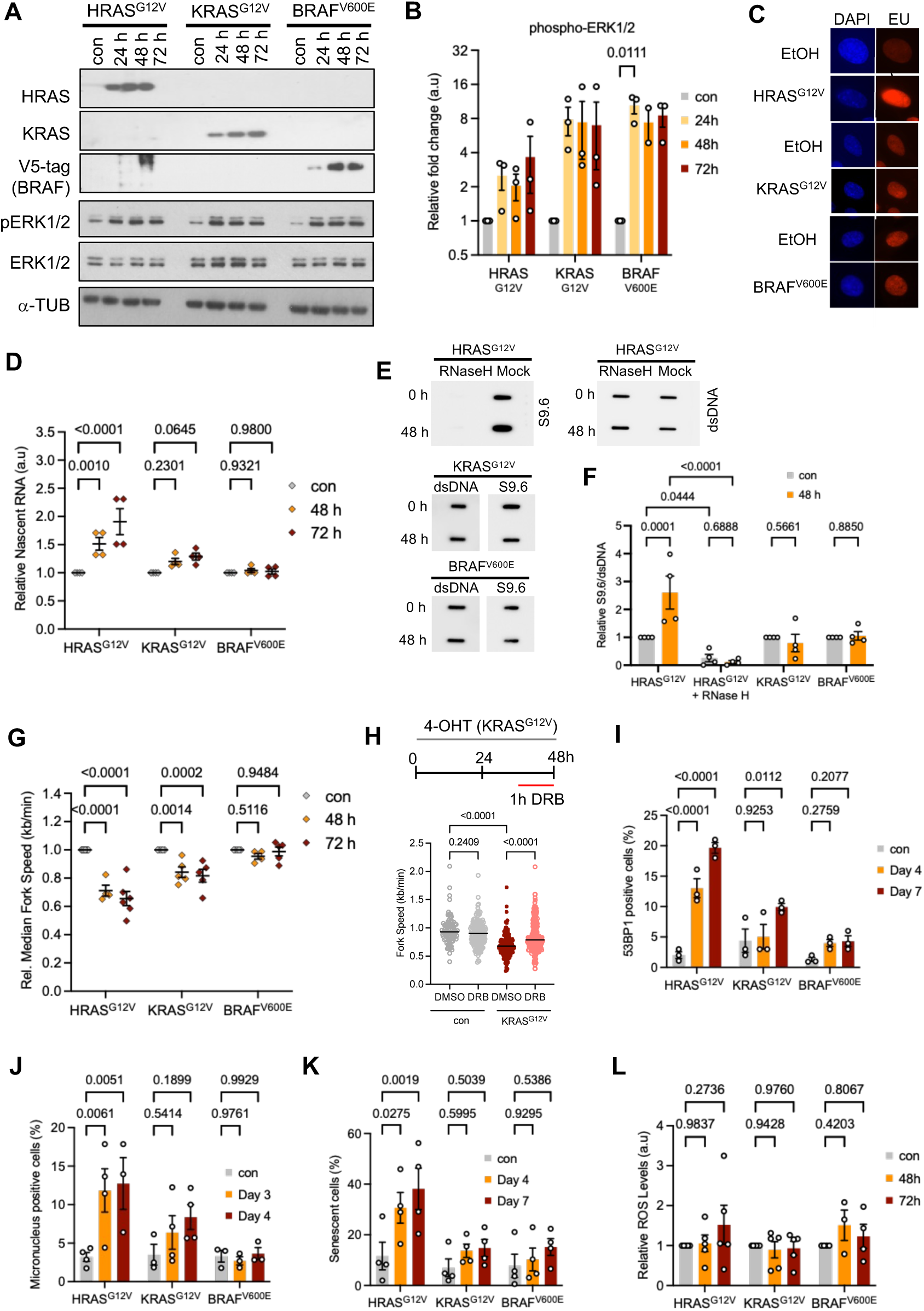
Hypertranscription and replication stress induced by oncogenic HRAS, KRAS and BRAF. (A) Protein levels of HRAS, KRAS, V5-tag (BRAF), pERK1/2, ERK1/2 and α-TUBULIN (loading control) in BJ-hTERT cells after oncogene induction for the times indicated. (B) Densitometry quantification of pERK1/2 levels, normalised to loading and control, after oncogene induction for the times indicated. N=3. (C) Nascent RNA synthesis after oncogene induction, measured by nuclear incorporation of EU (red) for 1 h. (D) Nuclear EU intensity after oncogene induction, normalised to control. N=4. (E) Slot blots of genomic DNA stained with S9.6 antibody (RNA:DNA hybrids) and double-stranded DNA (dsDNA; loading control) 48 h after oncogene induction. RNase H treatment was used to validate S9.6 antibody specificity. (F) RNA:DNA hybrid quantification as in E. N=4. (G) Relative median replication fork speeds after oncogene induction, normalised to control. (N=4-10). (H) Replication fork speeds after 48 h KRAS^G12V^ induction with DRB or DMSO treatment in the last 1 h. Data from 1 repeat. (I) Percentages of cells containing more than five 53BP1 foci after oncogene induction. N=3. (J) Representative images of cells with micronuclei and percentages of cells with micronuclei after RAS induction for 1-7 days. N=3-4. (K) Percentages of senescent cells, measured by β-galactosidase staining, after oncogene induction. N=4. (L) Relative levels of reactive oxygen species (ROS) after oncogene induction. N=3-6. Means +/-SEM (bars) are shown with 1-way or 2-way ANOVA or mixed effects analysis. Scatter graphs show median (line).

We next used DNA fibre assays to measure the effect of oncogene induction on replication fork progression (Fig. 1G, Fig. S1A). As reported before^8^, HRAS^G12V^ induction slowed down replication fork progression, suggesting high levels of replication stress. KRAS^G12V^ induction slowed replication fork progression to a lesser extent than HRAS^G12V^. BRAF^V600E^ induction caused no replication fork slowing. To further examine replication stress induced by KRAS^G12V^, we used transcription inhibitor 5,6-Dichloro-1-β-D-ribofuranosylbenzimidazole (DRB) to test the contribution of transcription-replication conflicts to replication fork slowing (Fig. 1H). DRB rescued KRAS^G12V^-induced fork slowing, as described for HRAS^G12V^ (ref^8^). We also generated a CaCo2 colon cancer cell line harbouring the tamoxifen-inducible KRAS^G12V^ construct (Fig. S1B), where KRAS^G12V^ induction had minimal effects on RNA synthesis and replication fork slowing (Fig S1C, D).

To investigate oncogene-induced DNA damage, we quantified nuclear foci of DNA damage marker 53BP1 and micronuclei. HRAS^G12V^ induced more DNA damage than KRAS^G12V^ or BRAF^V600E^ (Fig. 1I, J). As observed before in mouse embryonic fibroblasts^30^, HRAS^G12V^ induction resulted in higher levels of senescence and apoptosis than induction of KRAS^G12V^ or BRAF^V600E^ (Fig. 1K, Fig. S1E). Only HRAS^G12V^ activation led to a growth arrest within the first ten days (S1F-H).

These data suggest that mutant HRAS, but not KRAS or BRAF, cause high levels of replication stress, DNA damage and growth arrest due to different induction of hypertranscription. However, HRAS^G12V^ might also promote more production of reactive oxygen species (ROS)^31^, which could underlie replication stress, DNA damage and growth arrest. We therefore measured ROS levels using a flow cytometry reporter assay, which showed no differences in the levels of ROS induced by the three oncogenes (Fig. 1L).

These data suggest that despite its roles in proliferation signalling and cell cycle progression, MAPK signalling alone is insufficient to cause the phenotypes of RAS-induced hypertranscription and replication stress.

### Oncogenic HRAS activates a PI3K-AKT-GSK3β signalling axis

RAS activates additional signalling pathways other than MAPK, especially the PI3K-AKT pathway (Fig. 2A). Strikingly, Western blot of AKT phosphorylation showed that HRAS^G12V^ activated PI3K signalling much more strongly than KRAS^G12V^ or BRAF^V600E^ (Fig. 2B-D). This agrees with previous reports that, at the same protein expression level, HRAS^G12V^ activates PI3K signalling more strongly than KRAS^G12V^ (ref^32^).

**Figure 2.**
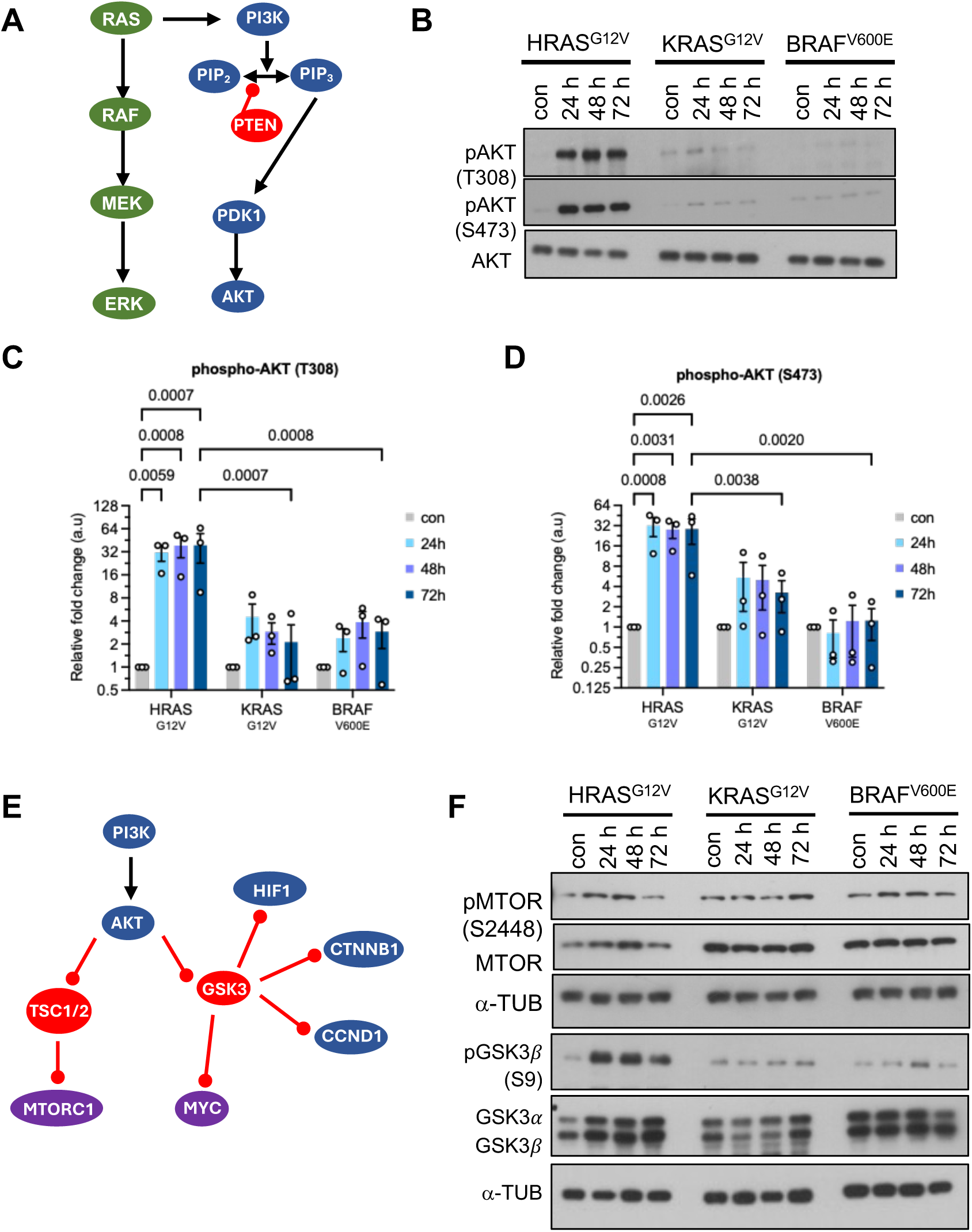
Oncogenic HRAS activates PI3K-AKT signalling. (A) Schematic of major signalling pathways downstream of RAS, BRAF and PI3K. PIP_2_: phosphatidylinositol-4,5-bisphosphate; PIP_3_: phosphatidylinositol-3,4,5-trisphosphate. (B) Protein levels of pAKT (T308, S473), and AKT after oncogene induction for the times indicated. (C) Densitometry quantification of pAKT (T308) levels, normalised to loading control, after oncogene induction for the times indicated. N=3. (D) Densitometry quantification of pAKT (S473) levels, normalised to loading control, after oncogene induction for the times indicated. N=3. (E) Schematic of major pathways for MTORC1 and MYV activation downstream of AKT. CTNNB1: β-Catenin; HIF1: Hypoxia-inducible factor 1. (F) Protein levels of pMTOR (S2448), MTOR, pGSK3 (S9), GSK3α, GSK3β, and α-TUBULIN (loading control) after oncogene induction. Means +/-SEM (bars) are shown with 2-way ANOVA.

PI3K signalling might stimulate RNA synthesis through activation of the MTORC1 or through inhibition of the GSK3 protein kinases, which in turn stabilises MYC and other proliferation factor such as CYCLIN D1^13, 25, 33^ (Fig. 2E). We therefore investigated the activation of MTORC1 signalling in response to oncogenic HRAS, KRAS or BRAF. Phosphorylation of MTOR or the MTORC1 downstream target RPS6 was comparable in response to HRAS^G12V^ and KRAS^G12V^ (Fig. 2F; Fig S2A). However, AKT-mediated phosphorylation of GSK3β was strongly increased in response to HRAS^G12V^ compared to KRAS^G12V^ or BRAF^V600E^ (Fig. 2F).

### Oncogenic HRAS increases E2F and MYC activity

To obtain further insight into HRAS versus KRAS-induced replication stress, we used steady-state RNA sequencing to obtain gene expression profiles. RNAseq was performed after 24h and 48h HRAS^G12V^ or KRAS^G12V^ induction (Fig. 3A, Supplementary Table S1). In agreement with the stronger oncogenic signalling, larger numbers of genes were significantly altered after HRAS^G12V^ induction compared to KRAS^G12V^ induction. In both cases, more genes were down-regulated than up-regulated (Fig. 3B). There was overlap between the genes upregulated by HRAS^G12V^ and KRAS^G12V^, but both oncogenes upregulated many unique genes (Fig. 3C). Functional enrichment analysis (biological process) showed that genes downregulated by both HRAS^G12V^ and KRAS^G12V^ were mostly involved in development, differentiation and cell structure and motility (Fig. 3D). Genes upregulated by HRAS^G12V^ were involved in ribosome biogenesis, translation, cell cycle and DNA replication, in agreement with the pro-proliferative effect of RAS signalling. Cell cycle and DNA replication were upregulated to a much smaller extent by KRAS^G12V^ and was detectable only if thresholds were lowered to 1.25-fold change across the analysis (Fig. 3E). We previously reported the potential importance of TBP up-regulation for RAS-induced hypertranscription and replication stress^8^, and TBP expression was only increased in response to HRAS^G12V^, but not KRAS^G12V^ (Fig. 3F).

**Figure 3.**
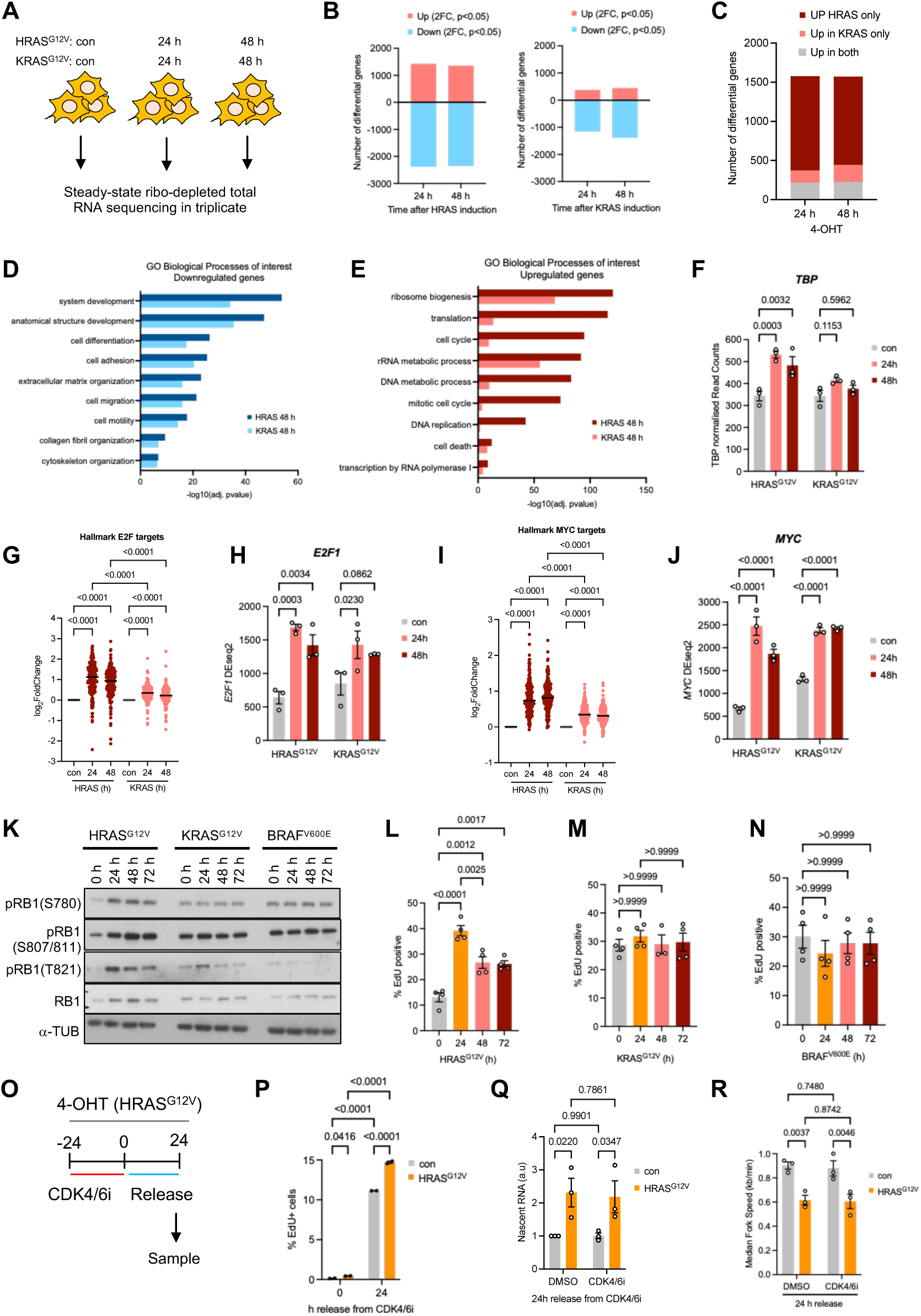
Oncogenic HRAS activates E2F and MYC. (A) Experimental setup for steady-state RNA sequencing. (B) Numbers of up- or downregulated genes (log2-fold change >1) after 24 or 48 h HRAS^G12V^ or KRAS^G12V^ induction. (C) Numbers of unique and shared up-regulated genes after HRAS^G12V^ or KRAS^G12V^ induction. (D) Functional enrichment analysis (gene ontology, biological process) of genes downregulated 48 h after HRAS^G12V^ or KRAS^G12V^ induction. (E) Functional enrichment analysis (gene ontology, biological process) of genes upregulated 48 h after HRAS^G12V^ or KRAS^G12V^ induction. (G) Log2 fold-change in hallmark E2F target gene expression after HRAS^G12V^ or KRAS^G12V^ induction. (H) *E2F1* expression (RNAseq, DEseq2) after HRAS^G12V^ or KRAS^G12V^ induction. N=3. (I) Log2 fold-change in hallmark MYC target gene expression after HRAS^G12V^ or KRAS^G12V^ induction. (J) *MYC* expression (RNAseq, DEseq2) after HRAS^G12V^ or KRAS^G12V^ induction. N=3. (K) Protein levels of pRB1 and α-TUBULIN after oncogene induction for the times indicated. (L) S phase percentage after HRAS^G12V^ induction as determined by EdU labelling and flow cytometry. N=4. (M) S phase percentage after KRAS^G12V^ induction. N=4. (N) S phase percentage after BRAF^V600E^ induction. N=4. (O) Experimental setup for CDK4/6 inhibitor (CDK4/6i) treatment and release. (P) S phase percentage after HRAS^G12V^ induction and treatment with CDK4/6i. N=2. (Q) Nuclear EU intensity after HRAS^G12V^ induction and release from CDK4/6i. N=3. (R) Average replication fork speeds after HRAS^G12V^ induction and release from CDK4/6i. N=3. Means +/-SEM (bars) are shown with 2-way ANOVA or mixed effects analysis.

It was previously shown that HRAS^G12V^ causes replication fork speeding rather than fork slowing in some immortalised human fibroblast lines^34^. This was linked to increased expression of topoisomerase 1 (TOP1) which reduces RNA:DNA hybrid levels^34^. However, TOP1 mRNA and protein levels were also increased after induction of HRAS^G12V^ and KRAS^G12V^ in BJ-hTERT fibroblasts (Fig. S3C, D). This suggests that RAS activation generally promotes TOP1 expression without this necessarily preventing replication stress.

We noticed that many of the HRAS^G12V^-upregulated genes were targets of the E2F family of transcription factors, which promote entry into and progression through S phase, or of MYC. HRAS^G12V^ induced expression of hallmark E2F target genes and *E2F1* (Fig. 3G, H) and of hallmark MYC target genes and *MYC* (Fig. 3I, J) more strongly than KRAS^G12V^. RT-qPCR showed that in contrast, BRAF^V600E^ did not increase *MYC* expression (Fig. S3E). This suggested that HRAS^G12V^ induction and increased PI3K signalling is associated with increased entry into S phase due to increased E2F activity.

### S phase entry is insufficient to cause replication stress

RAS signalling activates E2F through increased expression of D-type Cyclins and activation of CDK4 and CDK6, which in turn phosphorylate the retinoblastoma-associated protein (RB1), relieving the inhibition of E2F by RB1. Western blot analysis showed higher RB1 phosphorylation after HRAS^G12V^ induction compared to KRAS^G12V^ and BRAF^V600E^ induction (Fig. 3K). Flow cytometry to quantify 5-ethynyldeoxyuridine (EdU)-labelled replicating cells (Fig. S3F) confirmed increased S phase entry after HRAS^G12V^ induction (Fig. 3L). In contrast, there was no change in S phase content after KRAS^G12V^ or BRAF^V600E^ induction (Fig 3M, N).

Accelerated S phase entry might be an underlying cause for hypertranscription and replication stress. Nascent RNA synthesis has been reported to increase^35^ or decrease^36^ in S phase. EU labelling to quantify nascent RNA synthesis combined with CYCLIN A2 staining to assess cell cycle phase showed higher nascent RNA synthesis in S/G2 phase in control cells, but similar increases in RNA synthesis across G1 and S/G2 phases after HRAS^G12V^ induction (Fig S3G). Further to this, we previously found that down-regulating replication initiation with pan-CDK inhibitor roscovitine did not rescue HRAS^G12V^-induced replication fork slowing^8^. Nevertheless, we decided to use a small molecule inhibitor of CDK4/6, palbociclib, to specifically prevent S phase entry (Fig. 3O). CDK4/6 inhibition completely prevented S phase entry and reduced the difference in S phase content between control and HRAS^G12V^-induced cells after 24 h release from the drug (Fig. 3P). Nevertheless, CDK4/6 inhibition did not prevent hypertranscription (Fig. 3Q) or replication fork slowing (Fig. 3R). These data suggest that while increased E2F activity and S phase entry correlate with hypertranscription, they are not the cause of hypertranscription.

### HRAS stimulates nascent transcription of ribosome biogenesis genes, snoRNA synthesis and RNA Pol I activity

For an in-depth analysis of the impact of RAS signalling on ongoing transcription, we used sequencing of chromatin-bound RNA (chromatin RNAseq^37^) to quantify nascent transcripts after HRAS^G12V^ or KRAS^G12V^ induction for 48 h in triplicate (Fig. 4A; Fig. S4A, B; Supplementary Table S2). After principal component analysis, one repeat of KRAS^G12V^ induction was excluded as an outlier (Fig. S4B). Chromatin RNAseq showed more up-regulation of transcripts compared to steady-state RNAseq. Overall, HRAS^G12V^ up-regulated more transcripts than KRAS^G12V^ (Fig. 4B). Recently, nascent transcription was profiled in FUCCI-sorted BJ-hTERT cells^38^, allowing us to investigate RAS-induced changes by cell cycle phase. Both HRAS^G12V^ and KRAS^G12V^ downregulated genes which are mainly transcribed in G1 phase, while HRAS^G12V^ upregulated genes which are mainly transcribed in early G1 (EG1) and or late S/G2/M (G2) phases (Fig. 4C). HRAS^G12V^ increased transcription of more protein-coding genes, long non-coding RNAs (lncRNAs) and enhancer RNAs (eRNAs) (Fig. 4D-F). The relationship of the magnitude of transcriptional change with basal transcription activity or transcript length was similar in response to HRAS^G12V^ and KRAS^G12V^, with the largest RAS-induced changes observed in transcripts with low expression levels (Fig. S4C-F). In line with the increased S phase entry, histone genes were specifically upregulated by HRAS^G12V^ (Fig. 4G).

**Figure 4.**
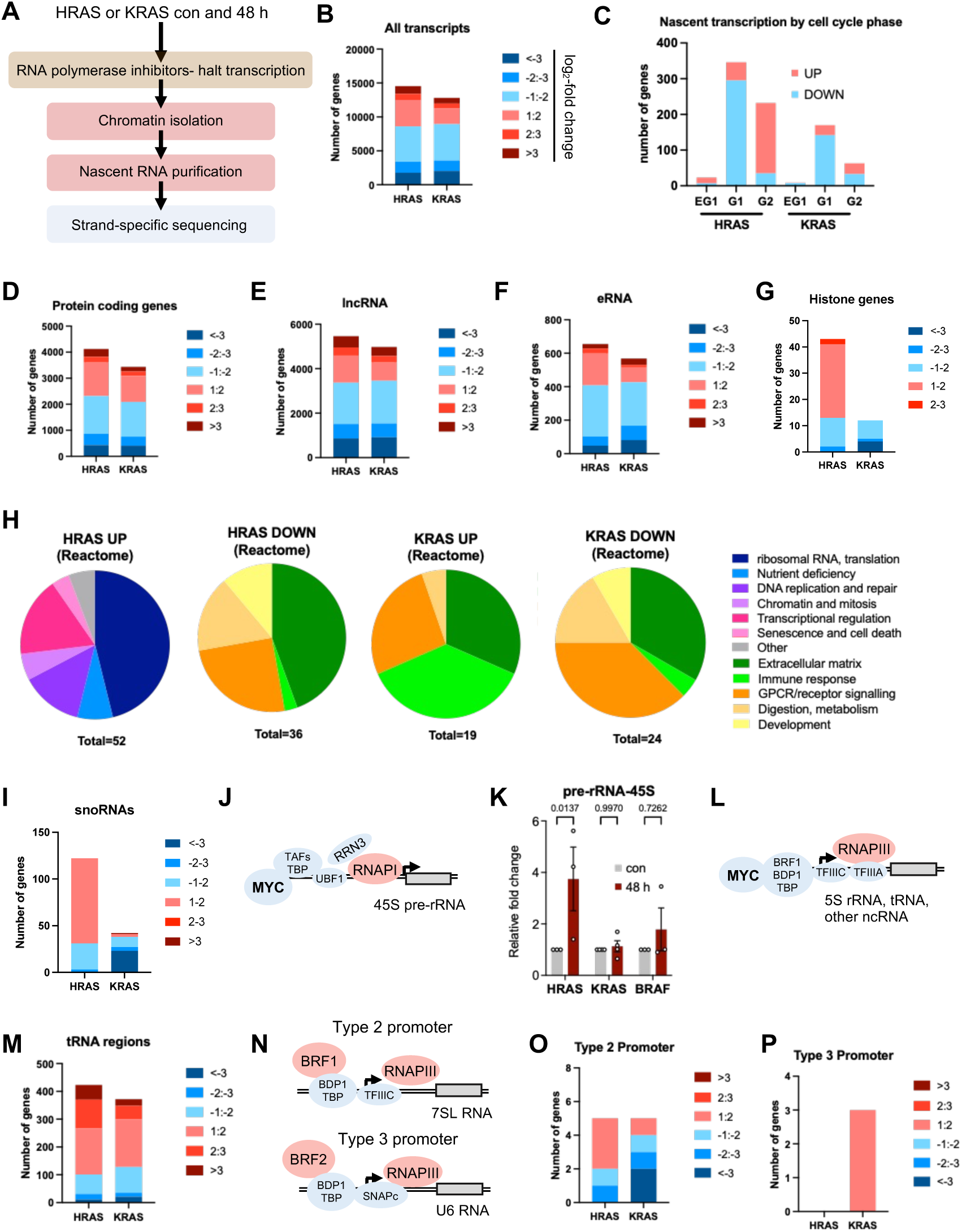
Oncogenic HRAS increases synthesis of transcripts involved in ribosome biogenesis and translation. (A) Principle of chromatin RNAseq approach. (B) Numbers of nascent transcripts up-or downregulated (log_2_-fold change) 48 h after HRAS^G12V^ or KRAS^G12V^ induction. HRAS^G12V^: N=3, KRAS^G12V^: N=2. (C) Numbers of nascent transcripts upregulated after HRAS^G12V^ or KRAS^G12V^ induction as in (B) that have been reported to be preferentially transcribed in either early G1 (EG1), G1 or late S phase, G2 and mitosis (G2)^38^. (D) Numbers of protein coding gene transcripts up- or downregulated 48 h after HRAS^G12V^ or KRAS^G12V^ induction. (E) Numbers of long noncoding (lncRNA) transcripts up- or downregulated 48 h after HRAS^G12V^ or KRAS^G12V^ induction. (F) Numbers of enhancer RNA (eRNA) transcripts up- or downregulated 48 h after HRAS^G12V^ or KRAS^G12V^ induction. (G) Numbers of histone gene transcripts up- or downregulated 48 h after HRAS^G12V^ or KRAS^G12V^ induction. (H) Functional enrichment analysis (REACTOME) of transcripts up- or downregulated 48 h after HRAS^G12V^ or KRAS^G12V^ induction. Numbers of significant REACTOME terms were counted. (I) Numbers of small nucleolar RNA (snoRNA) transcripts up- or downregulated 48 h after HRAS^G12V^ or KRAS^G12V^ induction. (J) RNA polymerase I (RNAPI) is activated by oncogene signalling to generate 45S pre-ribosomal RNA for ribosome biogenesis. (K) RT-qPCR analysis of chromatin-associated 45S pre-RNA transcripts after 48 h oncogene induction. N=3-4. (L) RNA polymerase III (RNAPIII) is activated by oncogene signalling to generate non-coding RNAs for ribosome biogenesis and translation. (M) Numbers of transcripts in tRNA regions up- or downregulated 48 h after HRAS^G12V^ or KRAS^G12V^ induction. (N) Components of RNAPIII type 2 and type 3 promoters. (O) Numbers of RNA polymerase III transcripts with type 2 promoters up- or downregulated 48 h after HRAS^G12V^ or KRAS^G12V^ induction. (P) Numbers of RNAPIII transcripts with type 3 promoters up- or downregulated 48 h after HRAS^G12V^ or KRAS^G12V^ induction. Means +/-SEM (bars) are shown with 2-way ANOVA.

Functional enrichment analysis (Reactome) showed that HRAS^G12V^ upregulated genes involved in cell cycle and transcription regulation, ribosome biogenesis and translation. Genes downregulated by HRAS and both up- or downregulated by KRAS were involved in extracellular matrix, immune response, cell signalling, and metabolism (Fig. 4H, I; Supplementary Table S3). Direct comparisons of up-regulated genes (Reactome) in the steady-state versus chromatin RNAseq datasets is shown in Fig. S4G, H.

In line with increased transcription of ribosome biogenesis genes, snoRNAs were specifically upregulated by HRAS. Increased expression of ribosome biogenesis genes and snoRNAs are consistent with increased MYC activity as snoRNAs are MYC targets^39^ (Fig. 4I). We then analysed changes in nascent transcription by RNA Pol I, which is activated by MYC to synthesise the 45S pre-ribosomal RNA (pre-rRNA)^40^ (Fig. 4J). We isolated chromatin-bound RNA followed by RT-qPCR of 45S pre-rRNA. RNA Pol I activity increased in response to HRAS^G12V^, but not KRAS^G12V^ or BRAF^V600E^ (Fig. 4K). Similarly, RNA Pol III is activated by MYC to synthesise the 5S rRNA, transfer (RNA) and other small non-coding RNAs involved in controlling transcription and translation^2, 40^ (Fig. 4L). HRAS^G12V^ increased transcription of more tRNA genes than KRAS^G12V^, suggesting a stronger stimulation of RNA Pol III activity (Fig. 4M). RNA Pol III target genes have three promoter types, which are distinguished by the presence of different basal transcription factors, either BRF1 and TFIIIC at type 1 and 2 or BRF2 and SNAPc at type 3^41^ (Fig. 4N). Type 1 promoters are at the 5S rRNA genes, whose transcripts are removed before sequencing. Type 2 promoters control the genes encoding tRNA and other RNAs involved in ribosome function and protein synthesis, while genes with type 3 promoters tend to encode transcripts that regulate RNA Pol II activity or mRNA splicing^41^. HRAS^G12V^ stimulated the transcription of non-tRNA type 2 promoter genes (Fig. 4O), but not of type 3 promoter genes (Fig, 4P).

Together these data support an increased activation downstream of HRAS^G12V^ of signalling factors such as AKT and MYC, which promote activity of RNA Pol I and RNA Pol III for ribosome biogenesis through increased expression and phosphorylation of transcription factors such as RRN3^40, 42, 43^.

### Inhibition or activation of PI3K modulates replication stress

To investigate the role of PI3K signalling in replication stress, we used small molecule inhibitors of MEK (PD0325901) and PI3K (GDC-0941) and measured the effect on RNA synthesis and replication fork slowing induced by HRAS^G12V^. Because both MEK and PI3K inhibitors arrest cells in G1 phase of the cell cycle, we developed a protocol exposing cells to inhibitors during the first 24 h of HRAS^G12V^ induction followed by release for 24 or 48 h before measuring RNA synthesis and replication fork progression (Fig. 5A). If MEK or PI3K inhibitor were present during the first 24 h of HRAS^G12V^ induction, this suppressed either AKT or ERK phosphorylation for up to 24h after release from the inhibitors (Fig. 5B; Fig. S5A, B). Cells re-entered S phase after 24 h release from the inhibitors (Fig. 5C; Fig. S5C). At the same time, the increase in RNA synthesis and the reduction in replication fork speeds was prevented by inhibition of PI3K or MEK for at least 24 h after release (Fig. 5D, E). MTORC1 inhibition with rapamycin had limited effect on RNA synthesis or replication fork slowing induced by HRAS^G12V^ (Fig. S5D, E).

**Figure 5.**
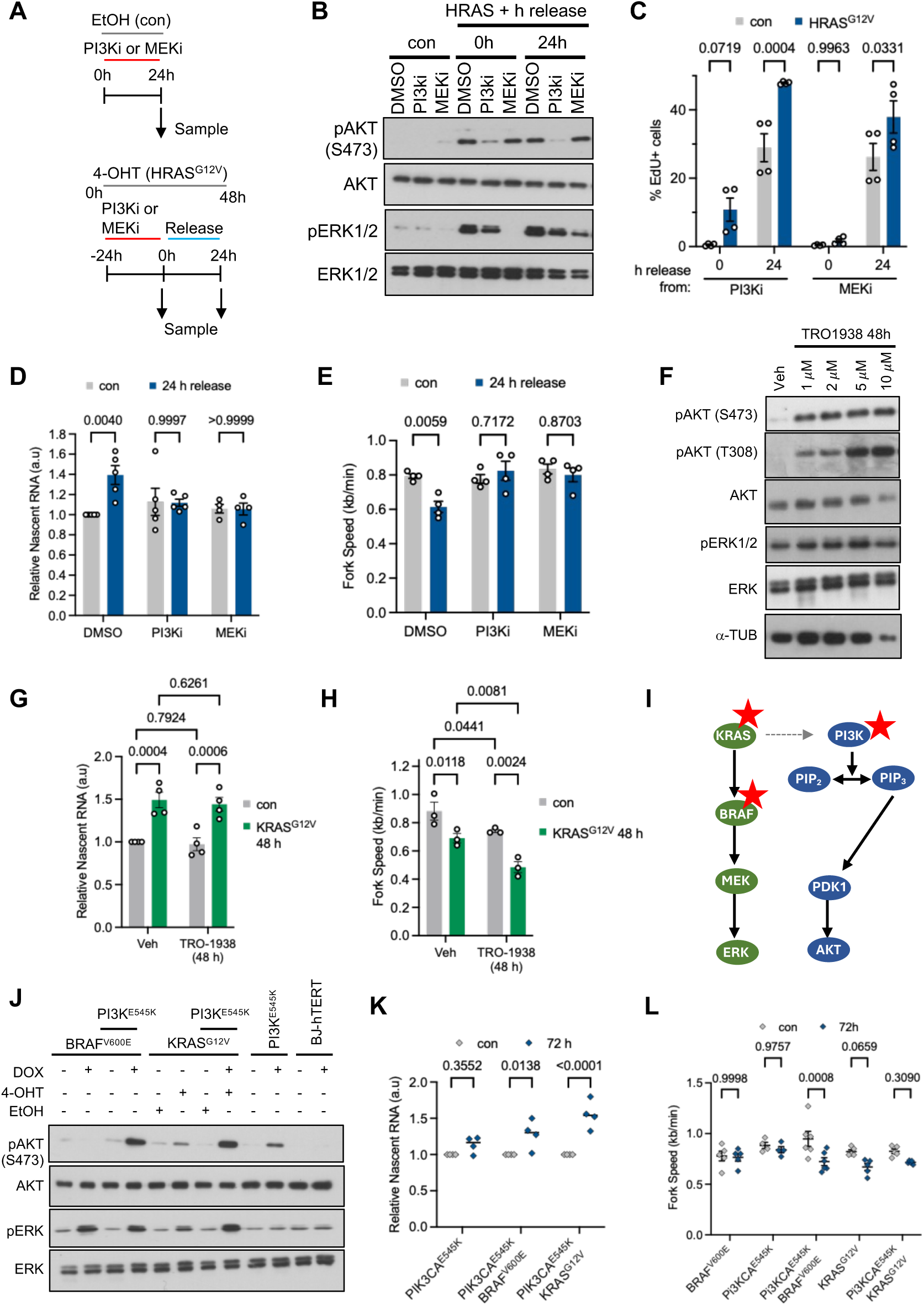
Inhibition or activation of PI3K modulates replication stress. (A) Experimental setup for PI3K inhibitor (PI3Ki) and MEK inhibitor (MEKi) treatment and release. (B) Protein levels of pERK1/2, ERK1/2, pAKT, and AKT after HRAS^G12V^ induction and treatment with MEKi or PI3Ki. (C) S phase percentage after HRAS^G12V^ induction with release from PI3Ki or MEKi as determined by EdU labelling and flow cytometry. N=4. (D) Nuclear EU intensity after HRAS^G12V^ induction and release from MEKi or PI3Ki. N=5 (MEKi N=4). (E) Average replication fork speeds after HRAS^G12V^ induction and release from MEKi or PI3Ki. N=4. (F) Protein levels of pAKT (S473), AKT, pERK1/2 and ERK1/2 after UCL-TRO-1938 treatment of uninduced BJ-hTERT-BRAF^V600E^ cells for 48 h. (G) Nuclear EU intensity after KRAS^G12V^ induction +/- UCL-TRO-1938 (5 μM) for 48 h. N=4. (H) Median replication fork speeds after KRAS^G12V^ induction +/- UCL-TRO-1938 (5 μM) for 48 h. N=3. (I) Inducible expression of oncogenic PI3K^E545K^, alone or with KRAS^G12V^ or BRAF^V600E^, was used to increase PI3K signalling, on its own or in combination with MAPK activation. (J) Protein levels of pAKT (S473), AKT, pERK1/2 and ERK1/2 after oncogene induction or doxycycline treatment (72 h) of parental cells (BJ-hTERT). (K) Nuclear EU intensity after oncogene induction. N=4. (EL) Median replication fork speeds after oncogene induction. N=4-6. Means +/-SEM (bars) are shown with 2-way ANOVA or mixed effects analysis.

We next attempted modelling the effect of PI3K activation alone or combined with RAS activation on hypertranscription and replication stress. For this we used the newly developed small molecule activator of PI3K, UCL-TRO-1938, which has been shown to stimulate both AKT and GSK3 phosphorylation^44^. UCL-TRO-1938 increased PI3K activity (Fig. 5F), and while it did not increase RNA synthesis any further (Fig. 5G), it caused replication fork slowing that was further exacerbated when combined with KRAS^G12V^ (Fig. 5H).

Finally, we generated cell lines for the inducible co-expression of oncogenic PI3K alone or in combination with either BRAF^V600E^ or KRAS^G12V^ to activate MAPK signalling (Fig. 5I). We successfully generated cell lines for inducible expression of PI3K with an activating mutation in the helical domain (PI3K^E545K^) but not in the kinase domain (PI3K^H1047R^). This suggests that the overexpression of the H1047R mutant may be detrimental to cells, possibly because it is a stronger activator of AKT^45^. The PI3K^E545K^ mutant nevertheless promoted AKT phosphorylation on S473, especially when combined with BRAF^V600E^ or KRAS^G12V^ (Fig. 5J). In contrast to UCL-TRO-1938, PI3K^E545K^ caused no additional replication stress when combined with KRAS^G12V^ compared to KRAS^G12V^ alone (Fig. 5K, L). However, combination of PI3K^E545K^ with BRAF^V600E^ increased nascent RNA synthesis and decreased replication fork speeds (Fig. 5K, L). PI3K^E545K^ did not increase S phase entry, whether alone or in combination with oncogenic BRAF or KRAS (Fig. S5F).

Altogether, PI3K hyperactivation can exacerbate hypertranscription and replication stress when combined with MAPK activation by oncogenic BRAF, but only small molecule activation of PI3K can increase replication stress alone or in combination with oncogenic KRAS, through a mechanism that may be separate from hypertranscription.

### PI3K and replication stress in cancer

We investigated the relationship of *BRAF*, *KRAS* and *PIK3CA* driver mutations with hypertranscription and replication stress markers in TCGA Pan Cancer Atlas datasets^46^. Several methods to estimate RNA output and therefore hypertranscription from cancer genomics datasets have been developed^17, 18^. Hypertranscription scores for TCGA Pan Cancer Atlas datasets are available from one of these studies^17^ (Fig. 6A), which also reported that transcription signatures of PI3K, AKT, MTORC1 and MYC activation correlated more strongly with hypertranscription than signatures of E2F or RAS activation^17^ (Fig. S6A). We then investigated co-mutations of *KRAS* or *BRAF* with *PIK3CA,* which occur with some frequency in colorectal adenocarcinoma (COAD) while *HRAS* mutations are rare in cancer. Analysis of COAD samples with available hypertranscription scores suggested that samples with combinations of *KRAS* or *BRAF* and *PIK3CA* mutation display a trend towards hyper transcription levels (Fig. 6B). We then employed a previously derived mRNA signature of oncogene-induced replication stress^47^. This signature was more strongly up-regulated by HRAS^G12V^ versus KRAS^G12V^ induction in our models (Fig. 6C, D). Analysis of this replication stress mRNA signature across COAD samples suggested that combinations of *KRAS* or *BRAF* with *PIK3CA* mutations displayed higher replication stress scores than samples with *KRAS* or *BRAF* mutations alone (Fig. 6E). Of note, many other factors may influence these results such as cancer subtype and p53 status.

**Figure 6.**
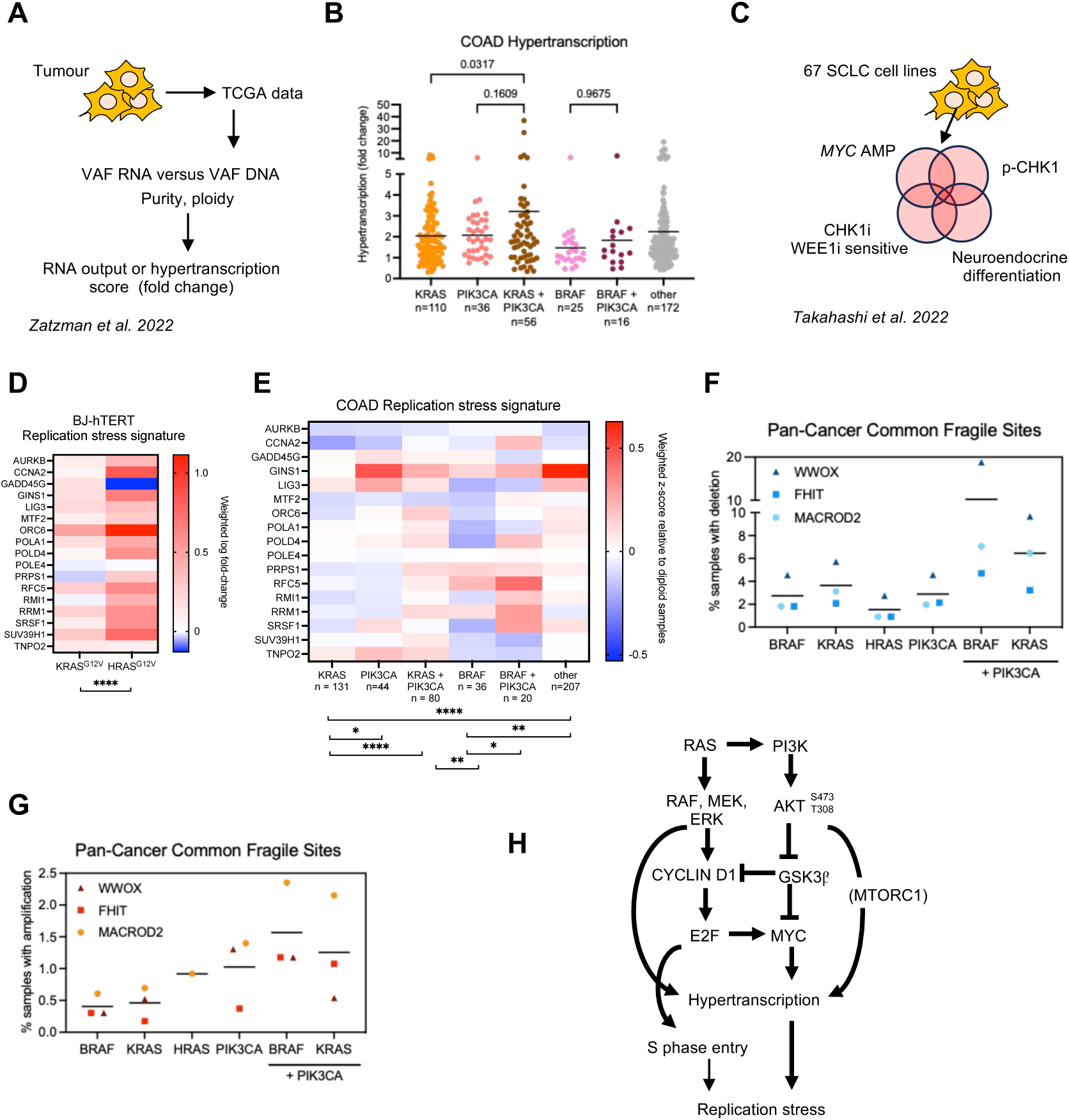
Markers of hypertranscription or replication stress in cancers with *PIK3CA* mutations. (A) Illustration of the approach used to generate hypertranscription scores^17^. (B) Hypertranscription scores (fold-change) in colorectal adenocarcinoma (COAD) samples with the indicated oncogene mutations. Lines denote mean. (D) Illustration of the approach to generate replication stress transcription signatures^47^. (E) Replication stress transcription signatures as in C after 48 h induction of KRAS^G12V^ or HRAS^G12V^ in BJ-hTERT cells (weighted log2-fold change). (E) Replication stress transcription signatures in COAD samples with the indicated oncogene mutations (weighted z-score). (F) Percentages of samples with homozygous deletions in the common fragile sites *WWOX*, *FHIT* or *MACROD2* across pan-cancer samples with the indicated oncogene mutations. (G) Percentages of samples with amplifications in the common fragile sites *WWOX*, *FHIT* or *MACROD2* across pan-cancer samples with the indicated oncogene mutations. (H) Model of the signalling pathways that promote TRCs downstream of RAS. PI3K and AKT contribute to S phase entry for example through GSK3β and CYCLIN D1, and to hypertranscription through AKT, MYC and potentially MTORC1. MAPK and PI3K pathways, and S phase entry and hypertranscription, cooperate in promoting TRCs and replication stress.

To analyse another readout of replication stress across cancer types, we used the PanCancer combined studies to quantify alterations in common fragile sites (CFS), which are known outcomes of replication stress. We quantified copy number alterations (CNAs) *WWOX*, *FHIT* and *MACROD2* the most commonly altered CFS in cancer^48^. Combinations of *HRAS* and *PIK3CA* mutations were too rare (27 out of 10,967 samples) to provide robust data. However, cancers with combinations of *KRAS* or *BRAF* and *PIK3CA* mutations had higher frequencies of CNAs in CFS than each mutation alone (Fig. 6F, G). These data support that PI3K pathway mutation can be associated with higher levels of replication stress than RAS/MAPK activation, and combined RAS/MAPK and PI3K mutation can further increase replication stress, especially in colon cancer where *KRAS* and *PIK3CA* are frequently co-mutated.

## DISCUSSION

Here we report that increased MAPK signalling is insufficient to promote RAS-induced hypertranscription and replication stress, at least in p53 wild type immortalised BJ fibroblasts, and that PI3K activation is an important driver of RAS-induced replication stress. Combined MAPK and PI3K pathway activation causes more replication stress than MAPK pathway activation alone. In the same cell background, HRAS^G12V^ is therefore a much more effective inducer of hypertranscription and replication stress than KRAS^G12V^ or BRAF^V600E^. Our findings suggest that AKT and ERK cooperate to induce replication stress by promoting hypertranscription and S phase entry, which they could promote either through direct signalling or via downstream activation of GSK3β, MYC, CYCLIN D and E2F (Fig. 6H).

Our findings shed much-needed light on replication stress induced by some of the most frequently activated oncogene signalling pathways. HRAS^G12V^ is a highly potent oncogene with the ability to cause different and more aggressive tumour types compared to KRAS^G12V^, or causing senescence when KRAS^G12V^ does not^30^. These differences resulted at least partly from more efficient protein expression of HRAS^G12V^ versus KRAS^G12V^ (ref^30^). However, HRAS^G12V^ activates PI3K signalling more strongly than KRAS^G12V^ even at the same level of protein expression^32^.

While our system uses protein stabilisation or overexpression to acutely activate RAS signalling, others have reported similar mild replication stress in cell lines with endogenous KRAS mutations compared to isogenic KRAS wild type counterparts^21^. Further in line with our findings, KRAS^G12V^ induction in RPE1-hTERT cells causes mild replication stress^49^ and BRAF^V600E^ induction in Caco2 cells only cause R-loop accumulation and replication stress after additional depletion of the RNA-binding protein SFPQ^50^.

PI3K^E545K^ alone did not cause replication stress phenotypes, which required combination with BRAF^V600E^. Similarly, hyperactivation of AKT in BJ-hTERT fibroblasts caused senescence but no DNA damage response^51^. However, our data suggest that combined PI3K-AKT and MAPK hyperactivation may predict hypertranscription and replication stress. Interestingly, low-level activation of BRAF and PI3K in cells harbouring doxycycline-inducible BRAF^V600E^ and PI3K^E545K^ (Fig. 5L) might increase fork speeds, an effect observed in several models of RAS activation^34, 52^. Chronic PI3K or AKT inhibition can cause replication stress^53, 54^. Here we show that chronic PI3K activation with UCL-TRO-1938 also causes replication stress, although it remains to be determined whether this involves transcriptional changes and TRCs. Our approaches for experimental PI3K activation could not recapitulate the very high levels of nascent RNA synthesis induced by HRAS^G12V^, which might require further changes such as increased S phase entry. The mechanisms of replication stress induction downstream of different RAS isoforms and PI3K mutants will require further investigation.

Both oncogenic HRAS and oncogenic KRAS hyper-activated the transcription of target genes of E2F, MYC and RNA Pol III. Much of this can be ascribed to comparable levels of MAPK and MTORC1 signalling, especially as ERK directly phosphorylates and activates key RNA Pol I and RNA Pol III transcription factors such as RRN3, UBTF and BRF1^55, 56, 57^. Post-transcriptional activation of RNA Pol III further requires MTORC1^58^.

Additionally, AKT signalling both through and independently of GSK3β and MYC is likely responsible for the qualitative difference between HRAS- and KRAS-induced replication stress as well as the inability of BRAF^V600E^ to cause overt replication stress. AKT can activate the RNA Pol I transcription factor RRN3 independently of MTORC1^42^. The contribution of the PI3K pathway to BRF1 activation deserves further investigation.

MYC could be an important effector of hypertranscription through AKT, as MYC can act as a “universal amplifier”, stimulating transcription by all three RNA polymerases^2, 59, 60^ and ribosomal biogenesis genes are MYC target genes^39,61^. We previously reported that TBP partially phenocopies HRAS^G12V^–induced replication stress^8^. MYC interacts with TBP and can promote TBP recruitment to promoters^62^.

GSK3β inactivation supports cell cycle progression through stabilising CYCLIN D1 and CYCLIN E1^33^, which might contribute to the increased S phase entry in cells with activated HRAS^G12V^. While our data suggest that S phase entry an hypertranscription can be uncoupled by MEK or PI3K inhibition (Fig. 2G-I), S phase entry may be required but not sufficient for RAS-induced hypertranscription and replication stress. S phase entry through increased E2F activity alone does not stimulate cell growth, which requires MYC^63^. Overexpression of Cyclin D1 or depletion of E2F inhibitor E2F6 increase rather than decrease replication fork speeds^64, 65^. However, strong replication stress was observed in hTERT-immortalised epithelial cells when overexpression of CYCLIN D1 and mutant CDK4 was combined with KRAS^G12V^ induction^49^.

PI3K pathway activation is common across cancer, with *PIK3CA* being the most frequently mutated oncogene in cancer^5, 6^. As such, the potential importance of PI3K for oncogene-induced replication stress has been somewhat neglected. Gene expression signatures of replication stress and hypertranscription scores have been developed to allow interrogation of cancer datasets^17, 18^. While the usefulness of such signatures may be limited by confounding factors such as proliferation or aneuploidy^66^, they nevertheless support that *PIK3CA*-mutant cancers can display signs of aberrant DNA replication and replication stress. Our findings further agree with the previously reported association between PI3K pathway activation and high hypertranscription scores in cancer datasets^17^.

Our findings highlight PI3K-AKT signalling as a previously underappreciated determinant of TRCs in RAS-driven cancers. In addition to PI3K-AKT activation, strong activation of E2F and MYC with high RNA Pol I activity and ribosome biogenesis gene expression could serve as markers for TRCs in RAS-mutant backgrounds. Understanding the role of PI3K-AKT signalling in promoting replication stress might aid optimal use of PI3K-AKT pathway inhibitors as well as therapeutic strategies targeting TRCs for personalised therapy.

## Methods

### Cell culture and cell line generation

Human BJ-hTERT HRAS^G12V-ER-TAM^ (ref^67, 68^), BJ-hTERT fibroblasts, RPE1-hTERT HRAS^G12V-TET-ON^ (ref^69^), HEK293FT (Thermo Fisher Scientific), and Caco-2 colorectal cancer cells were cultured in DMEM supplemented with 10% FBS, 1% L-glutamine and 1% penicillin-streptomycin at 37°C in 5% CO_2_ in a humidified incubator. All cell lines were authenticated using 17-locus short tandem repeat (STR) analysis (ATCC) and frequently tested for Mycoplasma infection.

Mutant BRAF (V600E) was synthesised by GenScript and cloned a doxycycline-inducible mutant pLIX403-ccdB-Blast (Addgene, Plasmid #158560) using gateway cloning. A doxycycline-inducible mutant PIK3CA^E545K^ expression vector was created by PCR subcloning of the PIK3CA^E545K^ coding sequence (Addgene, Plasmid #73055) into a modified pTRIPZ vector (Addgene, Plasmid #206981) lacking the TurboGFP coding sequence. KRAS^G12V^ vector was purchased from (Addgene, #35635). For lentiviral transduction, low-passage HEK293FT cells were transfected with lentiviral vectors contains the transgene of interest (See Supplementary Table 1), and the accessory plasmids VSV-G, RRE and REV (Addgene) for 24 h, using Lipofectamine 2000 (Invitrogen). Media containing the infectious virus was collected 24 h later, filter-sterilized, supplemented with 8 μg/ml polybrene and used to infect BJ-hTERT fibroblasts or Caco-2 cells for 24 h. This process was repeated 24 h later and cells were selected in either Puromycin, hygromycin or blasticidin for 3-7 days.

### Inhibitors

Small molecule inhibitors and activators were sourced as follows: MEK inhibitor PD0325901 (1 μM) and CDK4/6 inhibitor Palbociclib (1 μM) were obtained from Merck Life Science UK Limited. The PI3K inhibitor GDC-0941 (1 μM) and mTOR inhibitor Rapamycin (2.5 μM) were obtained from Stratech Scientific Ltd, while the PI3K activator UCL-TRO-1938 (5 μM) was sourced from Cambridge BioScience. The CDK9 inhibitor 5,6-Dichlorobenzimidazole riboside (DRB; 100 mM) was obtained from Sigma.

### DNA fibre analysis

Cells were sequentially labelled with 25 μM CldU and 250 μM IdU for 20 min each. Cells were harvested and resuspended in PBS at a concentration of 0.5-1 × 10^6^ cells per ml. 2 μl of cells spotting onto microscope slides and lysed with 7 μl of 0.5% SDS, 200 mM Tris-HCl pH 7.4 and 50 mM EDTA. Air-dried DNA fibre spreads were fixed in 3:1 methanol:acetic acid and stored at 4°C until required. DNA fibre spreads were rehydrated in H_2_O, denatured in 2.5 M HCl for 85 min, blocked with 1% BSA and 0.1% Tween-20 in PBS for 30 min and incubated with rat anti-BrdU (Abcam ab6326, 1:700) and mouse anti-BrdU (Becton Dickinson 347580, 1:500) for 1 h. Slides were then washed with PBS, fixed with 4% PFA and incubated with anti-rat AlexaFluor 594 (ThermoFisher, 1:500) and anti-mouse AlexaFluor 488 (ThermoFisher, 1:500) for 1.5 h. Fibres were examined using a Nikon E600 microscope with a Nikon Plan Apo 60x (1.3 NA) oil lens, a Andor Zyla sCMOS camera digital camera (C4742-95), and the NIS Elements BR 5.41.01 software (Nikon).

### Immunofluorescence

Cells grown on glass coverslips were fixed with 4% paraformaldehyde (PFA) for 10 min, permeabilised with 0.25% Triton X-100 for 5 min and blocked with 2% BSA and 0.05% Tween-20 in PBS. Coverslips were incubated with rabbit polyclonal anti-53BP1 (Bethyl A300-272A, 1:15,000) overnight at 4°C and anti-rabbit IgG AlexaFluor 594 (ThermoFisher A-11012, 1:500) for 1 h at room temperature. DNA was counterstained with 4′,6-diamidino-2-phenylindole (DAPI) and images were acquired on a Nikon E600 microscope with a Nikon Plan Apo 60x (1.3 NA) oil lens, a Andor Zyla sCMOS camera digital camera (C4742-95), and the NIS Elements BR 5.41.01 software (Nikon). Cells with more than 5-foci (53BP1) were “blind” scored as positive. Foci were quantified by eye directly on the microscope, and representative images were taken for illustration.

### 5-ethynyl uridine (EU) incorporation assays

Cells growing on glass coverslips were incubated with 1 mM EU in complete culture media for 1 h at 37°C in 5% CO_2_. Cells were then washed with PBS before fixing with 4% PFA for 15 min and permeabilised with 0.5% Triton X-100 for 15 min. The Click-iT reaction was performed using 4uM CuS0_4_, 100uM Ascorbic Acid and 0.15% Alexa Fluor® 594 in TBS for 30 min at room temperature. Cells were washed in TBS and mounted in DAPI. Imaging was performed immediately after Click reaction and nuclear masks (DAPI) were generated in ImageJ to quantify mean fluorescence intensities per nucleus. Results were normalized to control/DMSO to account for variation in staining intensity.

### SDS-PAGE and Western blotting

Whole cell extracts were obtained by sonicating cells in UTB buffer (50 mM Tris-HCl pH 7.5, 150 mM β-mercaptoethanol, 8 M urea). Protein concentration was determined by Bradford Assay (Bio-Rad) and proteins were denatured in Laemmli buffer at 80°C for 10 min. Polypeptides were separated by SDS-PAGE and transferred onto nitrocellulose membrane. Membranes were blocked for 1 h at room temperature using 5% milk or 5% BSA in TBS supplemented with 0.1% Tween-20 (TBS/T). Primary antibodies were diluted in blocking buffer overnight at 4 °C. Membranes were then washed with TBT/T followed by HRP-linked secondary antibody for 1 h at room temperature. A list of antibodies is provided in Supplemental Table 4. The signal was detected using ECL western blotting substrate (GE Healthcare).

### EdU, ROS and Annexin V flow cytometry

Cells were pulse-labelled with EdU for 30 min, followed by trypsinization and fixation in 4% PFA for 15 min. Permeabilization was carried out using 0.5% Triton X-100 in PBS for 20 min at room temperature. Subsequent EdU detection was performed via a Click-iT reaction utilizing CuSO_4_, ascorbic acid, and 0.15% Alexa Fluor® 488 in TBS (30 min, room temperature). The reaction was terminated by washing the samples in PBS containing 3% BSA. ROS detection using the Total Reactive Oxygen Species (ROS) Assay Kit 520nm (Invitrogen) and Annexin V staining (ThermoFisher Scientific) were performed according to the manufacturer’s instructions. Data were acquired on a BD LSR Fortessa X-20 and analysed using BD FACSDiva software.

### Cell proliferation and senescence assays

Cell proliferation was evaluated by seeding cells in 12-well plates at densities of 2.5-5x10^5^ cells per well. At specified time points, cultures were trypsinised and total cell number was counted using a haemocytometer. For senescence analysis, β-Galactosidase activity was detected using the Senescence β-Galactosidase Staining Kit (Cell Signaling), adhering strictly to the manufacturer’s protocol.

### S9.6 slot blot

Genomic DNA isolation was performed using the Qiagen DNeasy Blood & Tissue kit, following the standard protocol. For RNase H digestion, 10 µg of DNA per sample was incubated with 2 U/µg RNase H (NEB, M0297) or mock-treated for 2 h at 37°C. Subsequently, 250 ng of DNA was applied to a slot blot manifold and transferred to a pre-soaked Amersham Hybond N+ nylon membrane via vacuum filtration. Post-transfer, the membrane was UV-crosslinked (1200 µJ) and blocked for 1 h at room temperature in TBST containing 5% milk. Immunodetection was carried out by incubating the membrane overnight at 4°C with either mouse anti-DNA-RNA hybrid antibody (S9.6, 1:1000) or mouse anti-dsDNA (Abcam ab27156, 1:100,000) in 5% BSA/TBST. After washing with TBST, the blot was probed with HRP-linked goat anti-mouse secondary antibody (Cell Signaling 7074, 1:5000) in 5% milk/TBST for 1 h at room temperature. Signals were acquired using a BioRad ChemiDoc™ MP Imaging System and quantified with ImageJ.

### RNA extraction

All RNA extraction was performed using the Direct-zol RNA Miniprep Kit (Zymo Research) according to the manufacturer’s instructions. Briefly, cell or nuclei pellets were extracted in TRI Reagent® (Zymo Research) and stored at -80°C until required. RNA extraction was performed by adding equal volume of ethanol to TRI Reagent® before binding to bind directly to the Zymo-Spin™ Column. The column was washed according to the manufacturer’s instructions and incubated with DNase I to remove any DNA contamination. Purified RNA was eluted from the column using DNase/RNase-Free Water. RNA quality and concentration was determined using 4200 TapeStation (Agilent).

### RT-qPCR

cDNA was synthesised from 1 µg of the total RNA using qScript reverse transcriptase (Quanta Bioscience) using the following conditions: 22 °C, 5 min; 42 °C, 30 min; 85 °C 5 min; 4 °C. 2 µl of cDNA were analysed using a Real Time PCR QuantStudio 5 system with SYBR™ Green PCR Master Mix (ThermoFisher) and 1μM forward and reverse primers. Reactions encompassed initial denaturation for 10 min at 94°C, followed by 40 cycles for 10 sec at 94°C, primer-specific annealing for 30 sec, and 5 sec at 72°C. Primers and annealing conditions are included in Supplemental Table 5. Primer efficiency was calculated using absolute quantification from a standard curve.

### Steady state RNA sequencing

For steady state RNAseq messenger RNA was purified from total RNA using poly-T oligo-attached magnetic beads. After fragmentation, the first strand cDNA was synthesized using random hexamer primers followed by the second strand cDNA synthesis. The library was ready after end repair, A-tailing, adapter ligation, size selection, amplification, and purification. The library was checked with Qubit and real-time PCR for quantification and bioanalyzer for size distribution detection. Pooled quantified libraries were sequenced on Illumina platform NovaSeq 6000 S4.

### Chromatin-bound RNA (ChrRNA) sequencing

Cell fractionation was performed as previously described^37^ with modifications described below. All steps were conducted on ice or at 4°C and in the presence of 20 μM α-amanitin (Sigma, A2263), 25 μM RNA polymerase I inhibitor CX-5461 (Merck, 5.09265), 25 μM RNA polymerase III inhibitor ML-60218 (Merck, 557403), 50 Units SUPERaseIN (Life Technologies, AM2696) and Protease inhibitors cOmplete (Roche, 11873580001). Briefly, 5 × 10^6^ cells for each condition were resuspended and incubated in 0.15% NP-40, 10 mM Tris-HCl pH 7.0, 150 mM NaCl for 5min, then layered onto a sucrose buffer (10 mM Tris-HCl pH 7.0, 150 mM NaCl, 25% sucrose) and centrifuged at 16,000 g for 10 min. The supernatant carefully removed, and the nuclei pellet was gently resuspended in nuclei wash buffer (0.1% Triton X-100, 1 mM EDTA, in 1x PBS) and centrifuged at 1500 g for 1 min. The supernatant was removed, and the pellet was gently resuspended in glycerol buffer (20 mM Tris-HCl pH 8.0, 75 mM NaCl, 0.5 mM EDTA, 50% glycerol, 0.85 mM DTT) before an equal volume of nuclei lysis buffer (1% NP-40, 20 mM HEPES pH 7.5, 300 mM NaCl, 1M Urea, 0.2 mM EDTA, 1 mM DTT) was added. Samples were vortexed, incubated on ice for 2 min and centrifuged at 18,500 g for 2 min. The supernatant was removed, and the chromatin pellet was immediately incubated with TRI Reagent® (Zymo Research) and stored at -80°C for RNA extraction with Direct-zol RNA Miniprep Kit (see above).

Approximately, 200-300 ng of input RNA in each repeat were depleted of ribosomal RNA with NEBNext® rRNA Depletion Kit v2 according to manufacturer’s guidelines. Chromatin RNAseq was performed on chromatin fractions. Chromatin RNAseq libraries were prepared using NEBNext Ultra II 8482 Directional RNA Library Prep kit for Illumina (New England Biolabs) and all libraries were paired-end sequencing on NovaSeq 6000 (Illumina).

### RNAseq data analysis

Sequences were quality checked using FastQC(v0.11.9), (https://www.bioinformatics.babraham.ac.uk/projects/fastqc/) then quality filtered and trimmed using TrimGalore (v0.6.6). (https://www.bioinformatics.babraham.ac.uk/projects/trim_galore/). For RNAseq, genome mapping was performed with STAR (v2.7.10b)^70^ against the human reference genome (hg38 assembly). FeatureCounts (subread v2.03)^71^ was used to count reads and differential analysis was performed using DESeq2^72^, with *p* value calculated using the Wald test. Differential genes were identified by *P* value < 0.05 and LogFC>2.

### Chromatin-RNAseq data analysis

For Chromatin-RNASeq, quality checking, trimming and alignment was performed as above for RNASeq. Following alignment, samtools (v1.13)^70^ was used to retain only properly paired and mapped reads. Counts for genes and genomic regions were calculated using samtools^70^ bedcoverage with GENCODE v43 gene annotation^73^. Counts were also generated for specific gene lists: lncRNA (GENCODE v43 gene annotation), eRNA (BJ-fibroblast specific, EnhancerAtlas2.0)^74^, histone and snoRNA genes (based on gene symbols), tRNA (gtRNAdb 2.0)^75^, Type 2 and Type 3 promoter^41^, and satellite regions (RepeatMasker)^76^. RPKM and LogFC values were calculated using R Studio (v4.4.0)^77^, with differential genes identified by LogFC as indicated.

### Additional bioinformatic analysis

PCA plots were generated in RStudio using plot.PCR function. Gene ontology of up- and downregulated genes was performed using g:Profiler)^78^. Profile plots and heatmaps were generated using deepTools (v3.5.6)^79^.

Functional enrichment analysis was performed using g:Profiler^80^. The decision to display BP (biological process) or REACTOME in the figures was based on length of the results list. Human hallmark gene sets HALLMARK_MYC_TARGETS_V2 and HALLMARK_E2F_TARGETS were obtained from the Molecular Signatures Database (MSigDB)^81, 82^.

### TCGA data analysis

Cancer Genome Atlas dataset Colorectal Adenocarcinoma (COAD, TCGA, PanCancer Atlas), all complete samples (524 samples). Was accessed using cBioPortal^83^. ID numbers of samples with oncogenic mutations were obtained by searching for KRAS: MUT, BRAF: MUT, PIK3CA: MUT. Replication stress signatures were obtained from ref^47^. mRNA expression z-scores relative to diploid samples (RNA Seq V2 RSEM) were analysed using Microsoft Excel and GraphPad Prism 10. Weighting factors from ref^47^ were used to adjust z-scores. WWOX, FHIT and MACROD2 copy number alterations were obtained by searching TCGA PanCancer Atlas Studies (10,967 samples)^46^ for KRAS: MUT, BRAF: MUT, HRAS: MUT, PIK3CA: MUT.

### Statistical analysis

Bar graphs show the mean +/- s.e.m from independent biological replicates unless indicated otherwise. Numbers of independent biological repeats (N) are indicated in the figure legends. Statistical tests were performed using the GraphPad Prism 10 software, version 10.6.1 (799). Gaussian distribution was determined using GraphPad Prism. Paired or unpaired student’s *t*-test was used for two comparisons with Gaussian distribution or a Mann-Whitney test for two comparisons with non-Gaussian distribution. For multiple comparisons, one-or two-way ANOVA or mixed-effects analysis with Dunnett’s, Tukey’s or Sidak’s test for datasets with Gaussian distribution and Kruskal-Wallis test with Dunn’s test for datasets with non-Gaussian distribution.

## Data and Software Availability

Data from RNAseq experiments have been deposited under series accession number GSE322526 (GSE322524 as unique number for Chromatin-RNASeq; GSE322525 as unique number for RNASeq). The authors declare that all other data supporting the findings of this study are available within the article and its supplementary information files and from the corresponding authors upon reasonable request.

## Acknowledgements

R.D.W.K., C.W., A.B. and R.J.W. were supported by a Cancer Research UK Programme Foundation award to E.P. and A.K. (C25526/A28275). The E.P. lab was also supported by the Medical Research Council (MR/W00190X/1, MR/W031442/1). We thank Dr Gemma Regan-Mochrie and Prof Clare C. Davies (University of Birmingham) for the kind gift of the modified pTRIPZ-PIK3CA^E545K^ construct. We acknowledge the support of the University of Birmingham - Flow Cytometry Facility, RRID:SCR_027107, for providing access to equipment and technical expertise. We acknowledge the Genomics Birmingham Facility and Matthew Newbold at the University of Birmingham for support of chromatin RNAseq experiments.

## Author contributions

R.D.W.K. conceived, designed and performed experiments, analysed data and contributed to writing the paper; C. W. analysed data and contributed to writing the paper; C.T and R.J.W. designed and performed experiments and analysed data; A.K coordinated and supervised the project; E.P. conceived, coordinated and supervised the project, designed experiments, analysed data and wrote the paper.

## Competing financial interests

E.P. has a consulting contract with Storm Therapeutics. The other authors declare no competing financial interests.

## Notes

https://www.ncbi.nlm.nih.gov/gds/?term=GSE322526[Accession]

