## Supplemental data for "PI3K-AKT activation determines oncogenic RAS-induced hypertranscription and replication stress"

### Supplementary figures

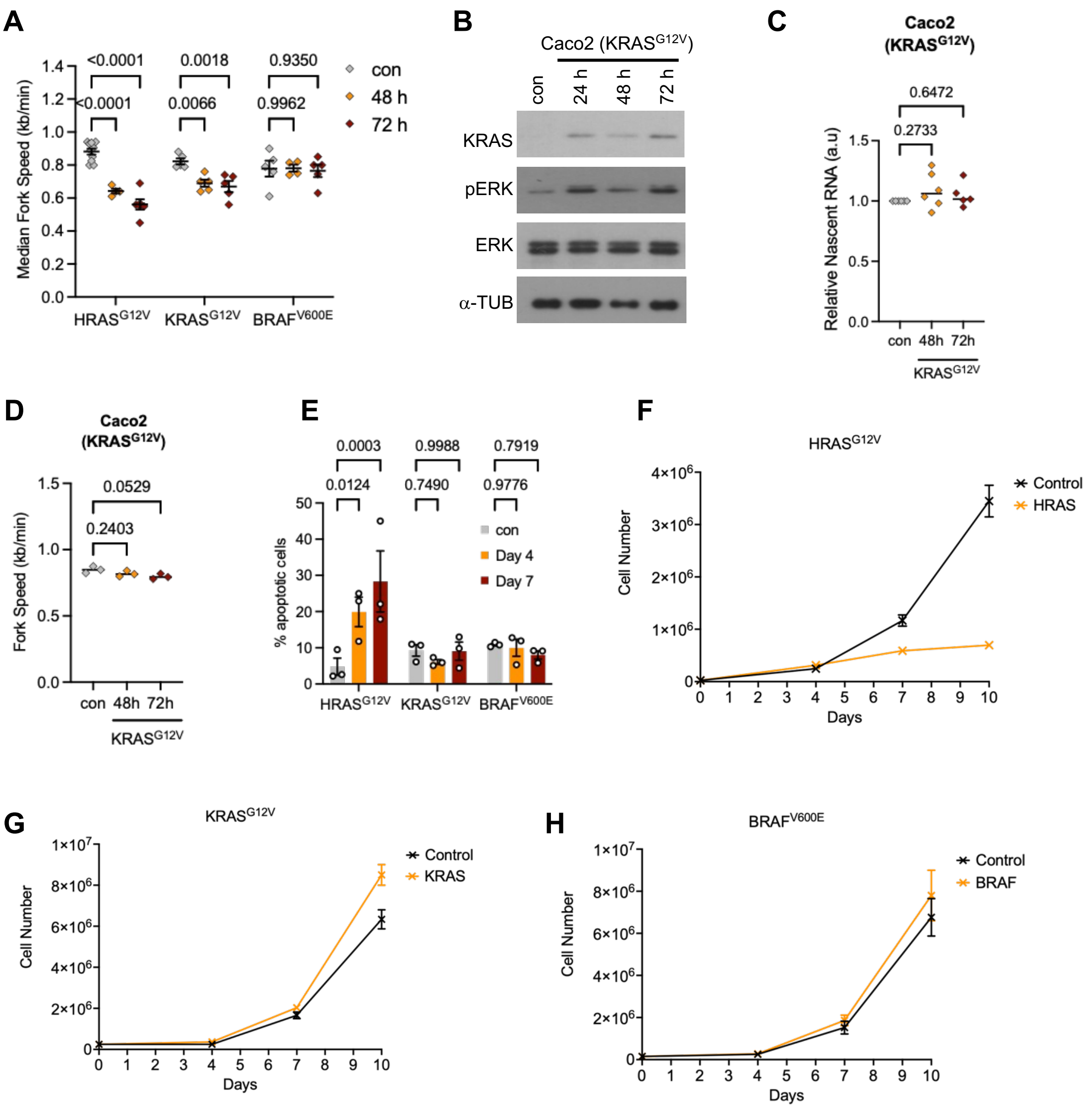


**Figure S1. Effect of induing oncogenic KRAS in CaCo2 cells and impact of oncogenic HRAS, KRAS and BRAF on apoptosis and cell growth**

(A) Median replication fork speeds after oncogene induction. (B) Protein levels of KRAS, V5-tag (BRAF), pERK1/2, ERK1/2 and α–TUBULIN (loading control) in CaCo2 cells after oncogene induction for the times indicated. (C) Nascent RNA synthesis after $KRAS^{G12V}$ induction in CaCo2 cells, measured by quantification of nuclear EU intensity. N=5-6. (D) Average replication fork speeds after KRAS induction in CaCo2 cells. N=3. E) Percentages of apoptotic cells, measured by Annexin V staining and flow cytometry, after oncogene induction in BJ-hTERT cells. N=3. (F) Growth curves of BJ-hTERT cells after $HRAS^{G12V}$ induction or EtOH control, measured by cell number. N=2. (G) Growth curves of BJ-hTERT cells after $KRAS^{G12V}$ induction or EtOH control, measured by cell number. N=2. (H) Growth curves of BJ-hTERT cells after $BRAF^{V600E}$ induction or no dox control, measured by cell number. N=2. Means +/-SEM (bars) with 1way or 2way ANOVA are shown.

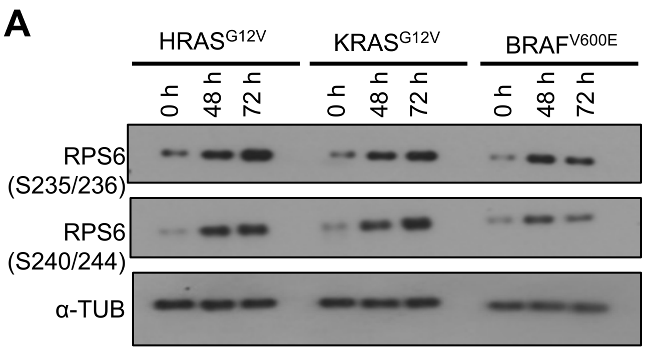


**Figure S2. MTORC1 activation downstream of oncogenic HRAS, KRAS or BRAF.**

(A) Protein levels of p-RPS6 (S235/236), p-RPS6 (S240/244) and α−TUBULIN (loading control) after oncogene induction for the times indicated.

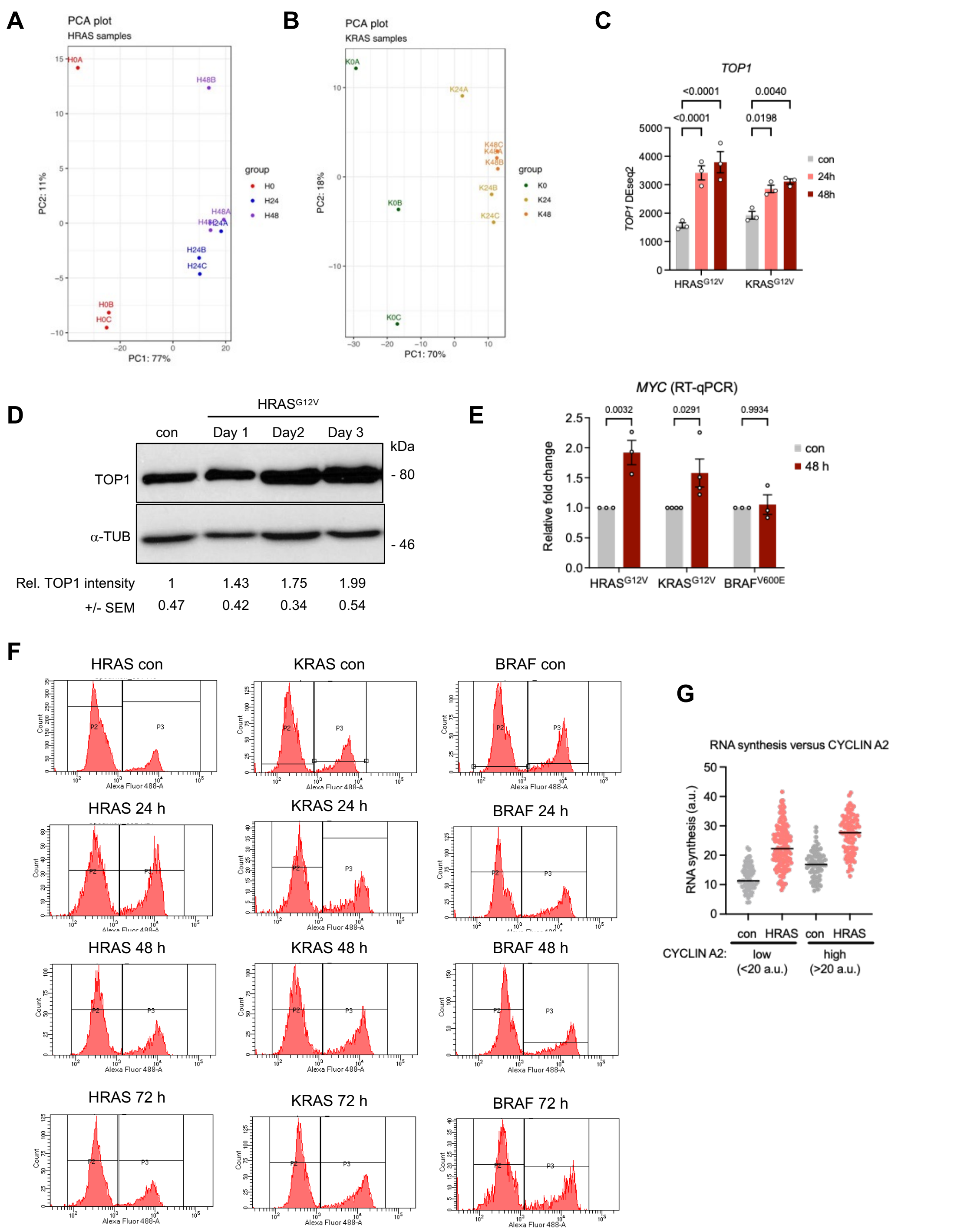


**Figure S3. Gene expression and cell cycle changes in response to oncogenic HRAS, KRAS or BRAF**

(A) Principal component analysis (PCA) plot of RNAseq samples from BJ-hTERT cells after EtOH control (H0) or 24 h (H24) or 48 h (H48) HRAS$^{G12V}$ induction. (B) PCA plot of RNAseq samples from BJ-hTERT cells after EtOH control (K0) or 24 h (K24) or 48 h (K48) KRAS$^{G12V}$ induction. (C) *TOP1* expression (RNAseq, DEseq2) after HRAS$^{G12V}$ or KRAS$^{G12V}$ induction. N=3. (D) Protein levels of TOP1 and α–TUBULIN (loading control) after EtOH control (con) or 1-3 days HRAS$^{G12V}$ induction. (E) RT-qPCR analysis of *MYC* expression in BJ-hTERT cells after oncogene induction. N=3 (KRAS: N=4).

(DF) Flow cytometry gating strategy for quantification of cell cycle distribution after oncogene induction. Cells were EdU-labelled and EdU-positive cells were detected by click reaction with AlexaFluor 488 (P3). (G) Nuclear EU intensity after $HRAS^{G12V}$ induction in cells with low (G1) or high (S/G2) CYCLIN A2 staining. N=1. Means +/-SEM (bars) are shown with 1way or 2way ANOVA.

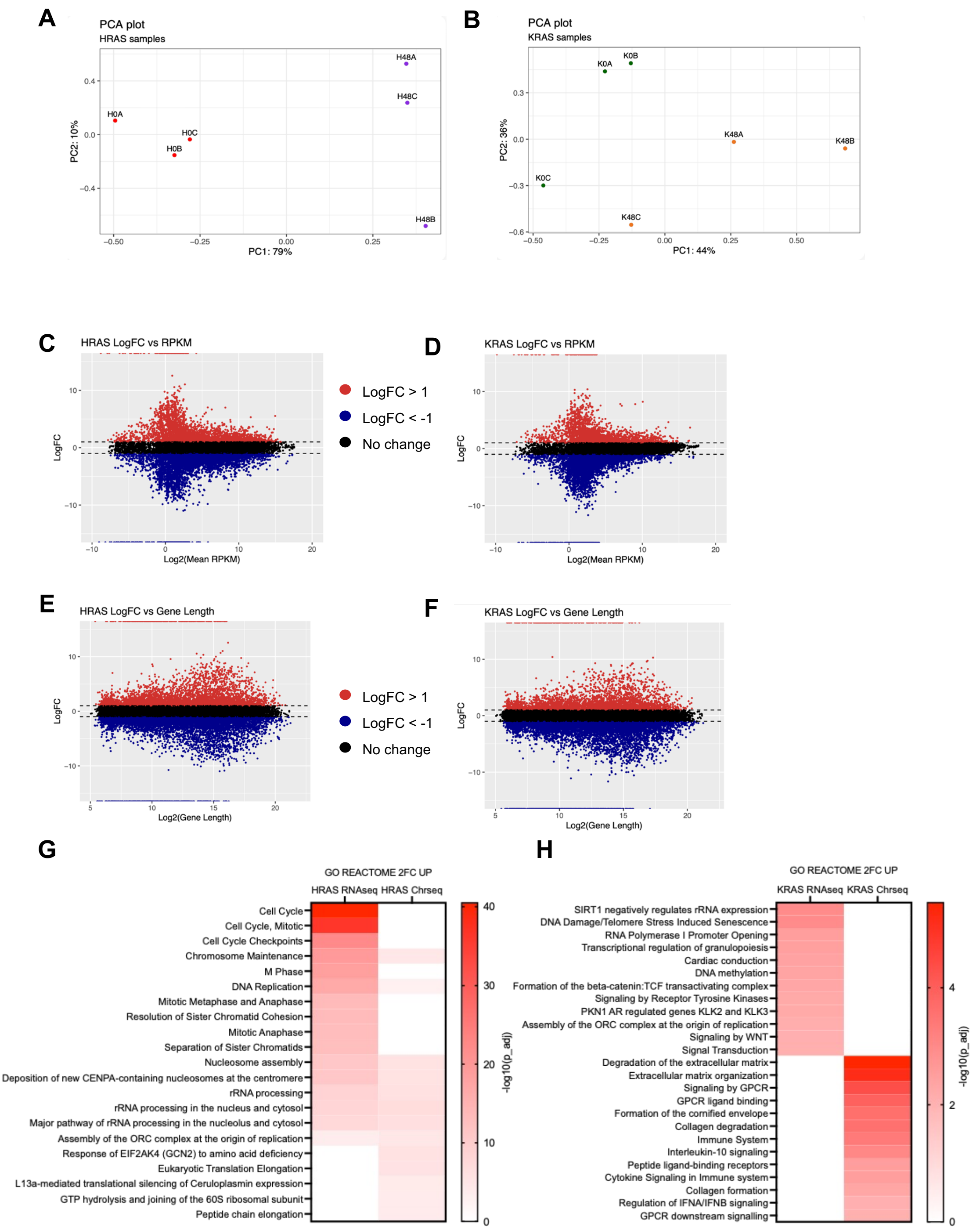


**Figure S4. Impact of oncogenic HRAS and KRAS on nascent transcription**
(A) Principal component analysis (PCA) plot of chromatin-RNAseq samples from BJ-hTERT cells after EtOH control (H0) or 48 h (H48) HRAS$^{G12V}$ induction. (B) PCA plot of chromatin-RNAseq samples from BJ-hTERT cells after EtOH control (K0) or 48 h (K48) KRAS$^{G12V}$ induction. Samples K0C and K48C were excluded from downstream analysis. (C) Expression change (log2-fold change) versus expression level (log2 of mean RPKM) in chromatin-RNAseq samples after HRAS$^{G12V}$ induction. (D) Expression change versus expression level in chromatin-RNAseq samples after KRAS$^{G12V}$ induction. (E) Expression change versus gene length (log2 of gene length) in chromatin-RNAseq samples after HRAS$^{G12V}$ induction. (F) Expression change versus gene length in chromatin-

RNAseq samples after KRAS$^{G12V}$ induction. (G) Functional enrichment analysis (g;Profiler, Reactome) of all genes upregulated 2-fold or more in RNAseq versus ChrRNAseq analysis after HRAS$^{G12V}$ induction for 48 h. Top 12 terms are shown. (H) Functional enrichment analysis (g;Profiler, Reactome) of all genes upregulated 2-fold or more in RNAseq versus ChrRNAseq analysis after HRAS$^{G12V}$ induction for 48 h. Top 12 terms are shown.

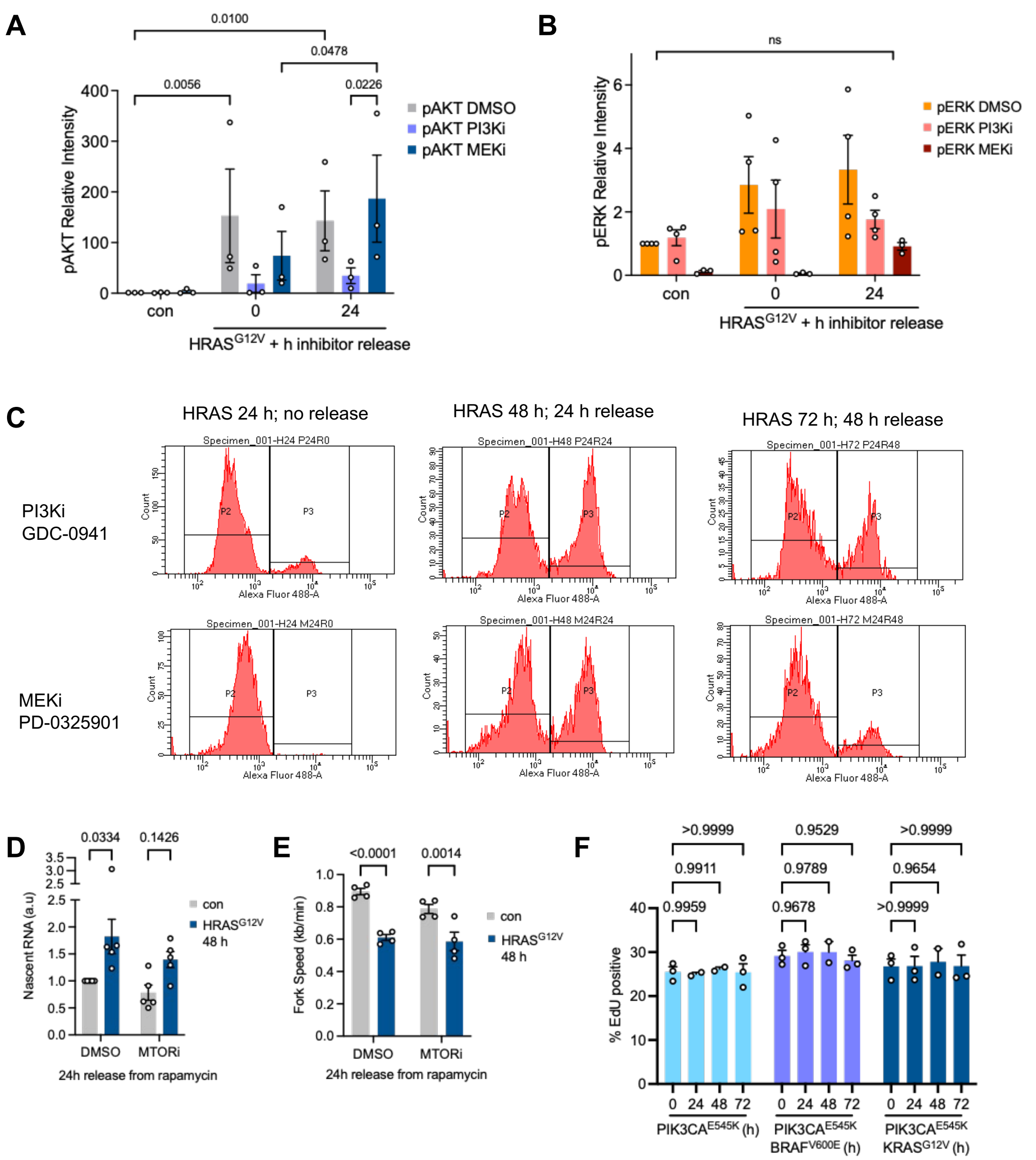


**Figure S5 Impact of PI3K, MEK and MTOR inhibition on downstream target phosphorylation, replication stress and cell cycle progression**

(A) Densitometry quantification of pAKT levels, normalised to loading control and con/DMSO, after EtOH control (con) or HRAS$^{G12V}$ induction for 24h plus 0 or 24 h release from PI3K or MEK inhibitor. N=3. (B) Densitometry quantification of pERK levels, normalised to loading control and con/DMSO, after EtOH control (con) or HRAS$^{G12V}$ induction for 24h plus 0 or 24 h release from PI3K or MEK inhibitor. N=3-4. (C) Flow cytometry gating strategy for quantification of cell cycle distribution after treatment with MEK or PI3K inhibitor. Cells were EdU-labelled and EdU-positive cells were detected by click reaction with AlexaFluor 488 (P3). (D) Nuclear EU intensity after HRAS$^{G12V}$ induction and release from Rapamycin (MTORi). N=5. (E) Average replication fork speeds after HRAS$^{G12V}$ induction and release from Rapamycin (MTORi). N=4. (F) S phase percentage after oncogene induction as determined by EdU labelling and flow cytometry. N = 3. Means +/-SEM (bars) are shown with 2way ANOVA or mixed-effects analysis are shown.

**A**

| Correlation with hypertranscription (TCGA pan-cancer) | |
|---|---|
| Rank (out of 50) | Hallmark gene set (GSEA-MSigDB) |
| 3 | MTORC1 SIGNALING |
| 7 | MITOTIC SPINDLE |
| 9 | PI3K AKT MTOR SIGNALING |
| 12 | MYC TARGETS V2 |
| 23 | MYC TARGETS V1 |
| 27 | G2M CHECKPOINT |
| 29 | DNA REPAIR |
| 30 | E2F TARGETS |
| 38 | KRAS SIGNALING DN |
| 50 | KRAS SIGNALING UP |

**Figure S6. PI3K pathway activation and hypertranscription in cancer.**
(A) Selected hallmark gene sets (GSEA-MSigDB) ranked by correlation of their expression with hypertranscription across cancer datasets, from Zatzman et al[17].

| Antibody | Supplier | Cat. No. | Species | Dilution |
|---|---|---|---|---|
| HRAS (F235) | Santa Cruz Biotechnology | sc-29 | Mouse | 1:500 |
| KRAS (11H35L14) | Invitrogen | 703345 | Rabbit | 1:2000 |
| V5 Tag | Invitrogen | R960-25 | Mouse | 1:500 |
| Akt (pan) mAb (C67E7) | Cell Signaling | 4691 | Rabbit | 1:2000 |
| Phospho-Akt (Ser473) mAb (D9E) | Cell Signaling | 4060 | Rabbit | 1:2000 |
| Phospho-Akt (Thr308) (D25E6) | Cell Signaling | 13038 | Rabbit | 1:1000 |
| p44/42 MAPK (Erk1/2) | Cell Signaling | 9102 | Rabbit | 1:2000 |
| Phospho-p44/42 MAPK (Erk1/2) (Thr202/Tyr204) | Cell Signaling | 9101 | Rabbit | 1:2000 |
| Phospho-mTOR (Ser2448) | Cell Signaling | 2971 | Rabbit | 1:1000 |
| mTOR | Cell Signaling | 2972 | Rabbit | 1:1000 |
| Phospho-Rb (Ser807/811) (D20B12) | Cell Signaling | 8516 | Rabbit | 1:1000 |
| Phospho-RB1 pThr821 | Invitrogen | 44-582G | Rabbit | 1:2000 |
| Phospho-Rb (Ser780) (D59B7) | Cell Signaling | 8180 | Rabbit | 1:1000 |
| Rb (4H1) | Cell Signaling | 9309 | Mouse | 1 in 1000 |
| Phospho-GSK-3 beta (Ser9) | Cell Signaling | 9323 | Rabbit | 1:1000 |
| GSK-3alpha/beta | Cell Signalling | 5676T | Rabbit | 1:5000 |
| Phospho-S6 Ribosomal Protein (Ser235/236) (D57.2.2E) | Cell Signaling | 4858 | Rabbit | 1:1000 |
| Phospho-S6 Ribosomal Protein (Ser240/244) (D68F8) | Cell Signaling | 5364 | Rabbit | 1:1000 |
| TOP1 | Abcam | ab85038 | Rabbit | 1:1000 |
| TUBA4A (TUBA1) | Sigma | T6074 | Mouse | 1:25000 |

**Table S4: List of antibodies used for Western blot**

| Primer | Sequence (5' - 3') |
|---|---|
| nPre-rRNA-45S-F1 | CCT GCT GTT CTC TCG CGC GTC CGA |
| nPre-rRNA-45S-R1 | AAC GCC TGA CAC GCA CGG CAC GGA |
| MYC | CAGCGACTCTGAGGAGGAAC |
| MYC | GCTGCGTAGTTGTGCTGATG |
| RPLP0 Forward (control) | CAGATTGGCTACCCAACTGTT |
| RPLP0 Reverse (control) | GGAAGGTGTAATCCGTCTCCAC |

**Table S5: Sequences for RT-qPCR primers**
